# CellUntangler: separating distinct biological signals in single-cell data with deep generative models

**DOI:** 10.1101/2025.01.10.632490

**Authors:** Sarah Chen, Aviv Regev, Anne Condon, Jiarui Ding

## Abstract

Single-cell RNA-seq data have provided new insights into intracellular and intercellular processes. Because multiple processes are active in each cell simultaneously, such as its cell type program, differentiation, the cell cycle, and environmental responses, their respective signals can confound one another, requiring methods that can separate and filter different complex biological signals. Each such signal is based on different gene activities and can define different relationships between cells. However, existing methods often focus on a single process or rely on overly restrictive assumptions, thus removing, rather than disentangling biological signals. Here, we develop CellUntangler, a deep generative model that embeds cells into a flexible latent space composed of multiple subspaces, each designed with an appropriate geometry to capture a distinct signal. We apply CellUntangler to datasets containing only cycling cells and both cycling and non-cycling cells, generating embeddings in which the cell cycle signal is disentangled from non-cell cycle specific signals, such as cell type or differentiation trajectory. We demonstrate CellUntangler’s extensibility by using it to capture and separate spatial from non-spatial signals. With CellUntangler, we can obtain latent embeddings that capture various biological signals and perform enhancement or filtering at the gene expression level for downstream analyses.

## Introduction

Single-cell RNA-seq (scRNA-seq) measurements of gene expression levels in individual cells have facilitated the identification of new cell types, the discovery of gene regulatory networks, and an enhanced understanding of spatial and temporal processes within and across cells^1^. Multiple processes are simultaneously active within each cell, including the cell cycle, cell differentiation, the cell’s type, and environmental responses. Each of these processes relies on different gene activities and the cells may relate to each other differently depending on the context (e.g., as separable clusters for cell types, continuous trajectories and bifurcations for differentiation, or a circle for the cell cycle, etc.).

This poses two analysis challenges. First, one or more prominent processes (e.g., cell cycle or cell type) may confound or obscure others during analysis, whereas an ideal analysis would uncover each of the processes and the cell relations in its contexts. Second, identifying and filtering out stronger signals (e.g., the cell cycle) to reveal any other residual ones is often a bespoke procedure that does not generalize well. Poor filtering of the effects from confounding factors can negatively impact downstream analysis of scRNA-seq data, but removal of a signal can also lead to loss of biological information. For example, consider the cell cycle. Removing cell cycle effects can uncover hidden signals, enabling the identification of distinct cell subpopulations^2^. Without proper removal, variability in gene expression among cells of the same type may be attributed to the cell cycle, causing cells to cluster by their cell cycle stage rather than their true biological identity^3^. In addition to being a confounder of biological signals, the cell cycle is itself a critical biological process and can differ between cell populations at different differentiation states^4^. Multiple methods have been developed for cell cycle inference and filtering. Some methods focus solely on removing the effect (e.g., ccRemover^3^) as a preprocessing step for downstream analyses, whereas others (e.g., tricycle^5^, DeepCycle^6^, reCAT^7^, and SC1CC^8^) infer cell cycle pseudotime but do not remove its effects. Methods that perform both inference and filtering often rely on assumptions that may limit their applicability. For instance, Cyclum^9^ embeds all cells onto a one-dimensional circle, which can incorrectly position non-cycling cells along the circle manifold. Moreover, while many existing methods are tailored exclusively to the cell cycle (or cyclic processes), other biological processes of interest remain largely unaddressed. For example, Cyclum is applicable only to cyclic processes; scPrisma^10^ can work for different signals and utilizes different topologies, but struggles with datasets that include non-cycling cells; SiFT^11^ requires multiple steps, including calculating the similarity between cells, and does not produce an embedding of cells for each signal of interest. All methods described usually require separate enhancing and filtering steps, followed by dimensionality reduction. Batch effect removal would be another step as well.

Here, we develop CellUntangler, a deep learning model trained end-to-end to output separate latent representations for different biological processes, while simultaneously addressing batch effects, eliminating the need for multiple, separate steps. CellUntangler leverages variational auto-encoders (VAEs)^12–14^, which have been shown to perform effectively in scRNA-seq analysis^15–19^. CellUntangler simultaneously captures and filters multiple biological signals. Unlike previous approaches, which typically use one latent space resulting in a single representation per cell, CellUntangler’s latent space is composed of multiple subspaces, each capturing a separate signal. The geometry of each subspace can be Euclidean or non-Euclidean^20, 21^, and users can select the geometry that best aligns with the nature of the signal of interest. It has previously been shown that non-Euclidean latent spaces can be beneficial for scRNA-seq data analysis^22, 23^. The CellUntangler decoder allows us to obtain data enhanced or filtered for a specific signal at the gene expression level, which is useful for downstream analyses.

We demonstrate CellUntangler’s effectiveness by applying it in several contexts. With a cycling cell line dataset and a mouse embryonic stem cells dataset, CellUntangler produced a latent representation that accurately captured the cell cycle signal and removed its confounding effects in a second latent representation. With a dataset of cycling and non-cycling immune cells, non-cycling cells clustered together in the latent subspace designed for the cell cycle signal, and the cell type identities of cycling cells were discernible in the latent subspace encoding cell cycle-independent information. With mouse pancreatic cells, CellUntangler separately captured cell cycle and differentiation trajectory signals in two non-Euclidean subspaces. Finally, in a mouse liver cell dataset, it successfully captured and separated the spatial zonation signal from other signals.

## Results

CellUntangler (**Fig. 1a**) takes as input an scRNA-seq dataset 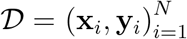 with *N* cells, where **x**_*i*_ is the unique molecular identifier (UMI) count vector of *G* genes in cell *i*, and **y**_*i*_ is a categorical vector indicating the batch or batches in which **x**_*i*_ is measured. We model **x**_*i*_ as being governed by a latent low-dimensional vector **z**_*i*_ composed of *k* distinct components, 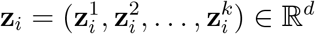. The final component, 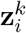 is used to capture additional signals not accounted for by the first *k*−1 latent components. For example, two components may be used: 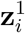 to capture the desired signal (e.g., cell cycle) and 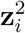 to capture signal-independent processes. To untangle the cell cycle or other desired signals into individual components 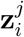, CellUntangler requires a set of marker genes for the target signal. Based on the marker genes for each component, CellUntangler decomposes **x**_*i*_ into *k* UMI count vectors, 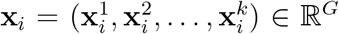 and uses a neural network decoder to model each UMI count vector 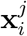 given **z**_*i*_. The batch vector **y**_*i*_ is also passed to the decoder to correct for batch effects. To estimate the posterior distributions for each component 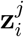, which are intractable to compute directly, we turn to variational inference and use a neural network encoder to output the parameters of the posterior distributions. We provide full details in the “Methods” section.

**Figure 1:**
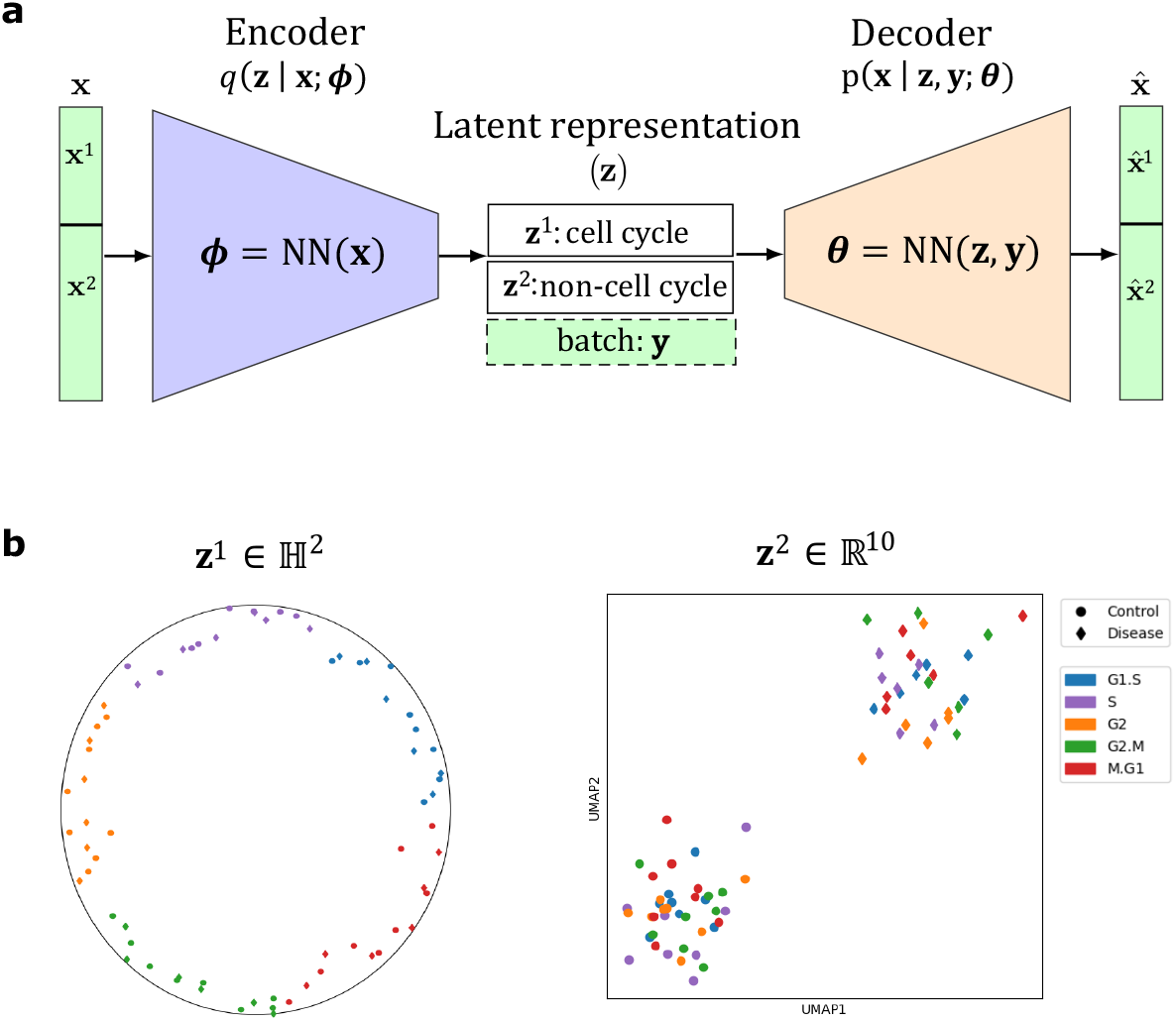
Method overview. CellUntangler takes as input scRNA-sequencing measurements and one or more batch effects, and embeds cells into decomposed latent subspaces with geometries tailored to separate distinct biological signals into individual representations. (**a**) The CellUntangler model with two latent subspaces: one to capture the cell cycle, **z**^1^, and the other to capture non-cell cycle-specific signals, **z**^2^. The parameters of *q*(**z** | **x**; ***ϕ***) and *p*(**x** | **z, y**; ***θ***) are represented by ***ϕ*** and ***θ***, respectively. (**b**) The cell cycle (cell cycle stage indicated by color) obscures the control and disease signal (indicated by shape). CellUntangler disentangles these signals by embedding the cell cycle signal in **z**^1^, revealing the control and disease signal in **z**^2^. **z**^2^ is projected to 2D using UMAP for visualization.

### CellUntangler captures and removes the cell cycle to reveal hidden biological signals

We initially focused on separating the cell cycle from other processes, as cycling cell profiles are expected to exhibit a circular structure (when considering cell cycle genes) due to the cyclical increase and decrease during cell cycle progression^24–26^. We modeled the latent distributions of the cell cycle in a hyperbolic space of dimension two, using the rotated hyperbolic wrapped normal distribution (RoWN)^27^ in the Lorentz model of the hyperbolic space. A hyperbolic space of dimension two provides a convenient embedding into a Poincaré disk when the Lorentz coordinates are projected to Poincaré coordinates, and the RoWN allows for local variation in the radial direction. We relied on a curated set of cell cycle marker genes (**Supplementary Table 1**). For cell cycle-independent processes, either the Euclidean or the hyperbolic space may be more suitable, depending on the nature of the signal. Importantly, using separate latent subspaces allows us to capture and remove the cell cycle signal from other signals (**Fig. 1b**).

We initially applied CellUntangler to scRNA-seq of wild type (WT) and Ago2 knockout (KO) cells^28^, showing that it effectively captured signals in two spaces: One (**z**^1^) reflecting the cell cycle (and mixing the genotypes) (**Fig. 2a**) and the other (**z**^2^) reflecting the genotype (and removing the cell cycle) (**Fig. 2b**). The cell cycle representation was in the hyperbolic space with the RoWN, **z**^1^ ∈ ℍ^2^, and the cell cycle independent one was in Euclidean space, **z**^2^ ∈ ℝ^10^. In **z**^1^ (**Fig. 2a**), cells also followed the correct progression of the cell cycle stages: G1.S → S → G2 → G2.M → M.G1, and back to G1.S. Conversely, a standard scRNA-seq processing and visualization pipeline^29–34^, mixed the two signals (**Fig. 2c**), with the cell cycle dominating over the genotype.

**Figure 2:**
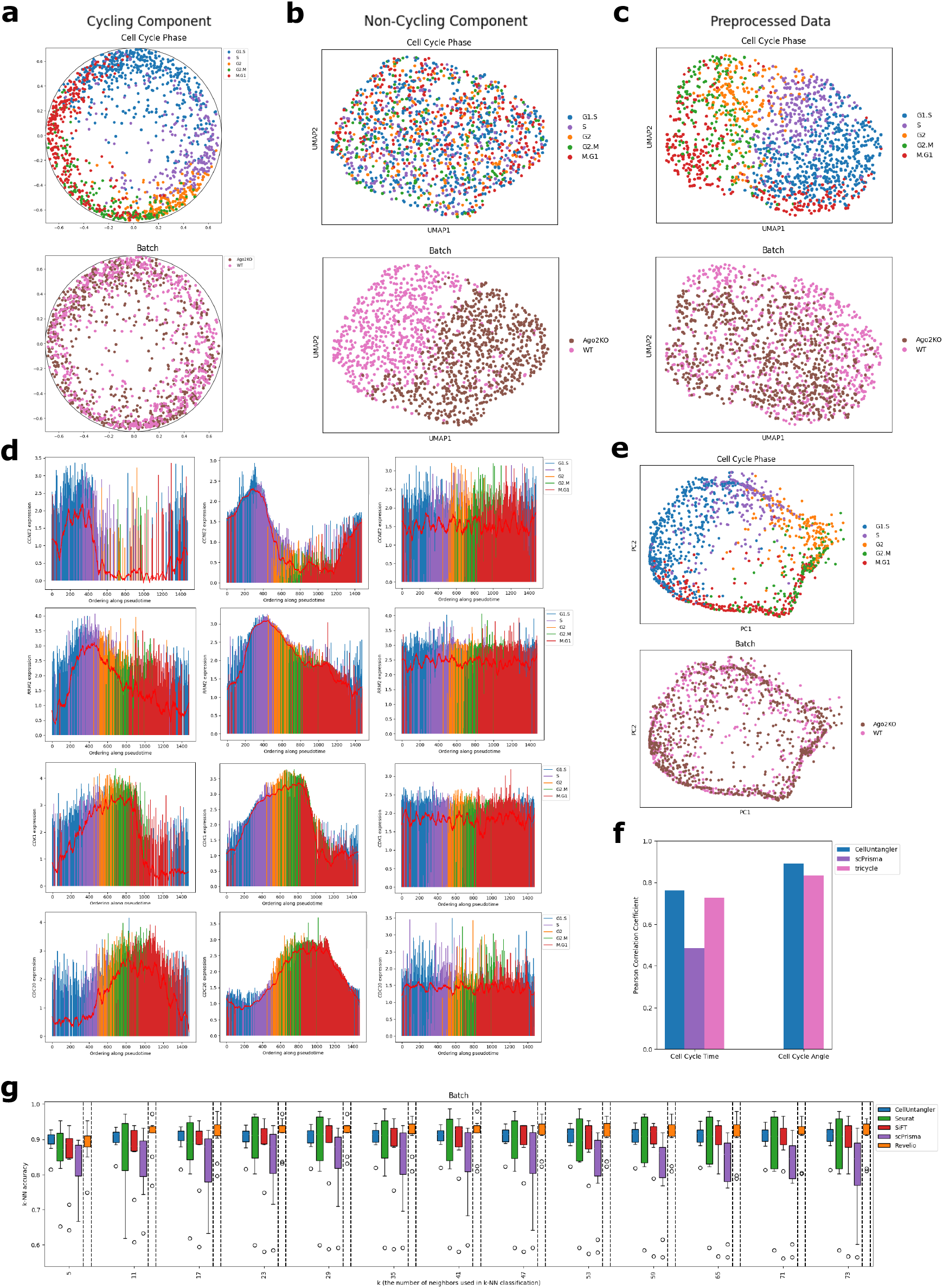
CellUntangler allows us to separate the cell cycle signal from wild type (WT) and knockout (Ago2KO) signal present in the HeLa dataset. (**a**) The first component of CellUntangler, **z**^1^, captures the cell cycle. (**b**) The second component, **z**^2^, projected to 2D using UMAP, filters out the cell cycle and separates wild type cells and knockout cells. (**c**) When using the standard preprocessing pipeline, cells separate by cell cycle stage. Wild type and knockout cells are mixed together. (**d**) Gene expression of *CCNE2* (G1.S), *RRM2* (S), *CDK1* (G2), and *CDC20* (M) as a function of pseudotime obtained with the embeddings from CellUntangler using the original preprocessed data (size-factor normalized and log-transformed, left) and the reconstructed gene expression output from CellUntangler after decoding **z**^1^ (middle) and **z**^2^ (right). Cell cycle stages (G1.S, S, G2, M) are in parentheses. (**e**) PCA representation of the reconstructed data obtained by running **z**^1^ through CellUntangler’s decoder. (**f**) Comparison of cell cycle reconstruction. (**g**) Accuracy of *k* -nearest neighbors on wild type and knockout cells using **z**^2^ from CellUntangler and different methods. Boxplots depict the medians and the interquartile ranges (IQRs). The whiskers are the lowest datum still within 1.5 IQR of the lower quartile and the highest datum still within 1.5 IQR of the upper quartile. Individual points below and above the whiskers indicate outliers.

We validated the cell cycle signal by examining the expression of individual genes in cells ordered along a pseudotime, obtained by projecting **z**^1^ from Lorentz to Poincaré coordinates, then measuring the angle of the projected Poincaré coordinates for each cell relative to the origin of the Poincaré disk, (0, 0) (**Fig. 2a**; Methods) (**Fig. 2d**, left and **Supplementary Fig. 1a**, left). Cell cycle-related genes exhibited expected expression peaks. For example, *CCNE2* peaked shortly after the G1.S transition; *RRM2* peaked during S phase, followed by *CDK1* peaking at the G2.M transition; and *CDC20* peaked at M phase (**Fig. 2d**, left).

By passing **z**^1^ to CellUntangler’s decoder, we enhanced the cell cycle signal present in the data at the gene expression level, yielding smoother gene expression trends with reduced noise (**Fig. 2d**, middle). This enhancement was consistent across other possible cell cycle marker genes used by Schwabe *et al*.^28^ to capture the cell cycle signal present in the same dataset (**Supplementary Fig. 1a**, middle), even though these genes were not provided as marker genes for our model.

Similarly, passing **z**^2^ to CellUntangler’s decoder filtered out the cell cycle signal at the gene expression level. Expression of cell cycle-related genes was flattened (**Fig. 2d**, right and **Supplementary Fig. 1a**, right). Performing PCA on the enhanced and filtered data further confirmed the effectiveness of CellUntangler: the cell cycle signal was either enhanced or filtered out as intended (**Fig. 2e** and **Supplementary Fig. 1b**), and vice versa for the genotype.

The quality of our cell cycle reconstruction compared favorably compared to other methods (**Fig. 2f** ; Methods). To assess the effectiveness of cell cycle removal, we used *k*-nearest neighbors (*k* -NN) classification accuracy (**Fig. 2g**) to distinguish between knockout and wild type cells, using 10-fold cross-validation. CellUntangler was among the top-performing methods for smaller values of *k*. Although Revelio performed better, it relies on a larger number of marker genes and requires prior knowledge of the association between genes and specific cell cycle phases (Methods).

### CellUntangler captures and removes the cell cycle in mouse embryonic stem cells

We next applied CellUntangler to a mouse embryonic stem cells dataset (using all available genes), originally presented by Riba *et al*.^6^ to study the cell cycle in this context. We used the hyperbolic space with the RoWN, **z**^1^ ∈ ℍ^2^, to capture the cell cycle and the Euclidean space, **z**^2^ ∈ ℝ^10^, to capture non-cell cycle-specific signals. The results from CellUntangler agreed with the expression phase from the original study, as the **z**^1^ vectors corresponding to the cells formed a circle, where the initial expression phase, *θ* = 0.0, progressed to the final expression phase, *θ* = 1.0 (**Fig. 3a**). Additionally, the cell cycle effects were non-discernible in **z**^2^ (**Fig. 3b**).

**Figure 3:**
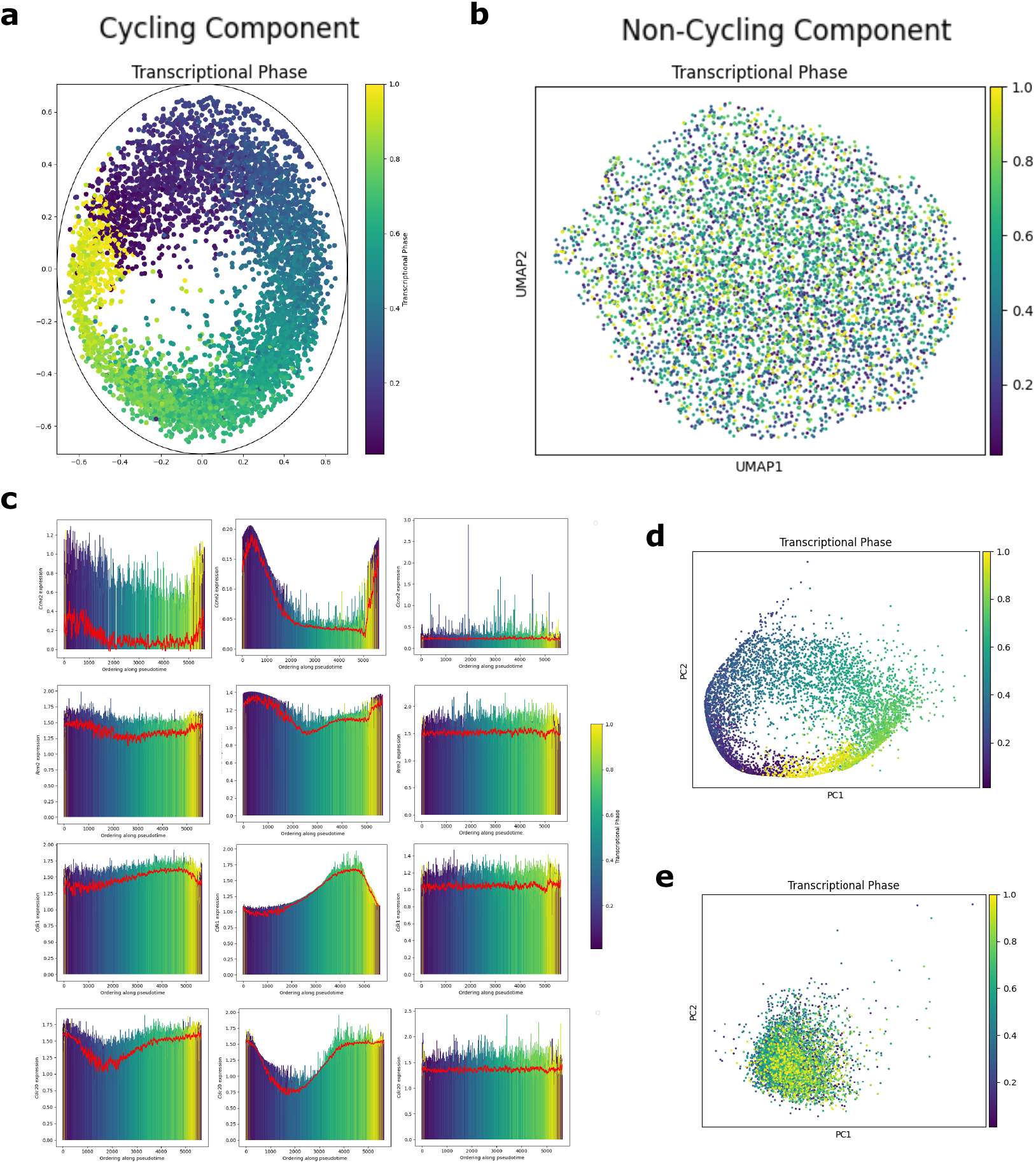
Analysis of the embeddings from CellUntangler on the mESCs demonstrates the capturing and removal of the cell cycle signal. (**a**) Visualization of the embeddings for the first component, **z**^1^, where cells were colored by the transcriptional phase output from the original study, and (**b**) embeddings for the second component **z**^2^. (**c**) Marker gene expression across cells when ordered by pseudotime obtained using **z**^1^ and colored using the transcriptional phase from DeepCycle of the original preprocessed data (size-factor normalized and log-transformed, left), and the reconstructed gene expression output from CellUntangler after decoding **z**^1^ (middle) and **z**^2^ (right). From top to bottom, *Ccne2* (G1.S), *Rrm2* (S), *Cdk1* (G2), and *Cdc20* (M). (**d, e**) PCA representations of the data reconstructions obtained from **z**^1^ (**d**) and **z**^2^ (**e**).

As above, we validated the results by examining the expression of cell cycle genes in cells ordered by pseudotime (**Fig. 3c**, left and **Supplementary Fig. 2a**, left), calculated by measuring the angular position of cells relative to the origin in **z**^1^ (**Fig. 3a**; G1 at *θ* = 0.0 to M at *θ* = 1.0, Methods). Expression patterns follow expected trends for *Ccne2* (peaks at G1.S transition), *Rrm2* (peaks at S phase) and *Cdk1* (peaks at G2.M transition), and *Cdc20* (peaks at M phase). CellUntangler’s data reconstructions based on **z**^1^ showed reduced expression noise in cell cycle genes, while the reconstruction from **z**^2^ exhibited flattened expression in these genes (**Fig. 3c**, middle and left). The effect on other cell cycle marker genes^28^, not included in the input list was similar (**Supplementary Fig. 2a**, middle and right). The cell cycle signal was enhanced in the PCA representation of the data reconstruction obtained with **z**^1^, and effectively filtered on the data reconstruction with **z**^2^ (**Fig. 3d, e**).

### CellUntangler provides insight into the cell cycle in populations containing both cycling and non-cycling cells

ScRNA-seq datasets typically contain both cycling and non-cycling cells. Although the cycling cells may belong to different types or subsets, they will often all group together in analysis due to the prominent signal from one program (cell cycle) dominating others (cell type programs), making it difficult to determine their cell type identities. Conversely, non-cycling cells can make capturing the cell cycle more challenging, as methods may try to assign a phase to each cell assuming it is cycling, or rely on a topological prior, causing them to fail as they attempt to place all cells on a circular manifold. Datasets may also contain technical and biological batch effects, further complicating the analysis.

To assess the ability of CellUntangler to tackle such challenges, we analyzed a myeloid cell atlas collected across multiple tissues, donors (of both biological sexes), ages, and lab techniques, and previously annotated as classical and nonclassical monocytes, conventional type one dendritic cells (DC1s), conventional type two dendritic cells (DC2s), migratory dendritic cells (migDCs), MNP/T doublets, and cycling cells (without a cell type assignment)^35^. As above, we used a hyperbolic space with the RoWN, **z**^1^ ∈ ℍ^2^ to capture the cell cycle, and a Euclidean space, **z**^2^ ∈ ℝ^10^, to capture the non-cell cycle-specific information.

In the cycling component (**z**^1^), the cycling cells were initially placed along the edge of the Poincaré disk, while the other cells were distributed throughout the interior of the disk (**Fig. 4a**). Following recentering the origin of the disk, we obtained the expected circular arrangement of cycling cells ordered along the disk circumference in the expected sequential order (**Fig. 4b**, top), surrounding non-cycling cells (**Fig. 4b**; Methods). In both the original and recentered disks, non-cycling cells were clearly mixed with respect to cell type (**Fig. 4a**, bottom and **Fig. 4b**, bottom), confirming that **z**^1^ captured only cell cycle information.

**Figure 4:**
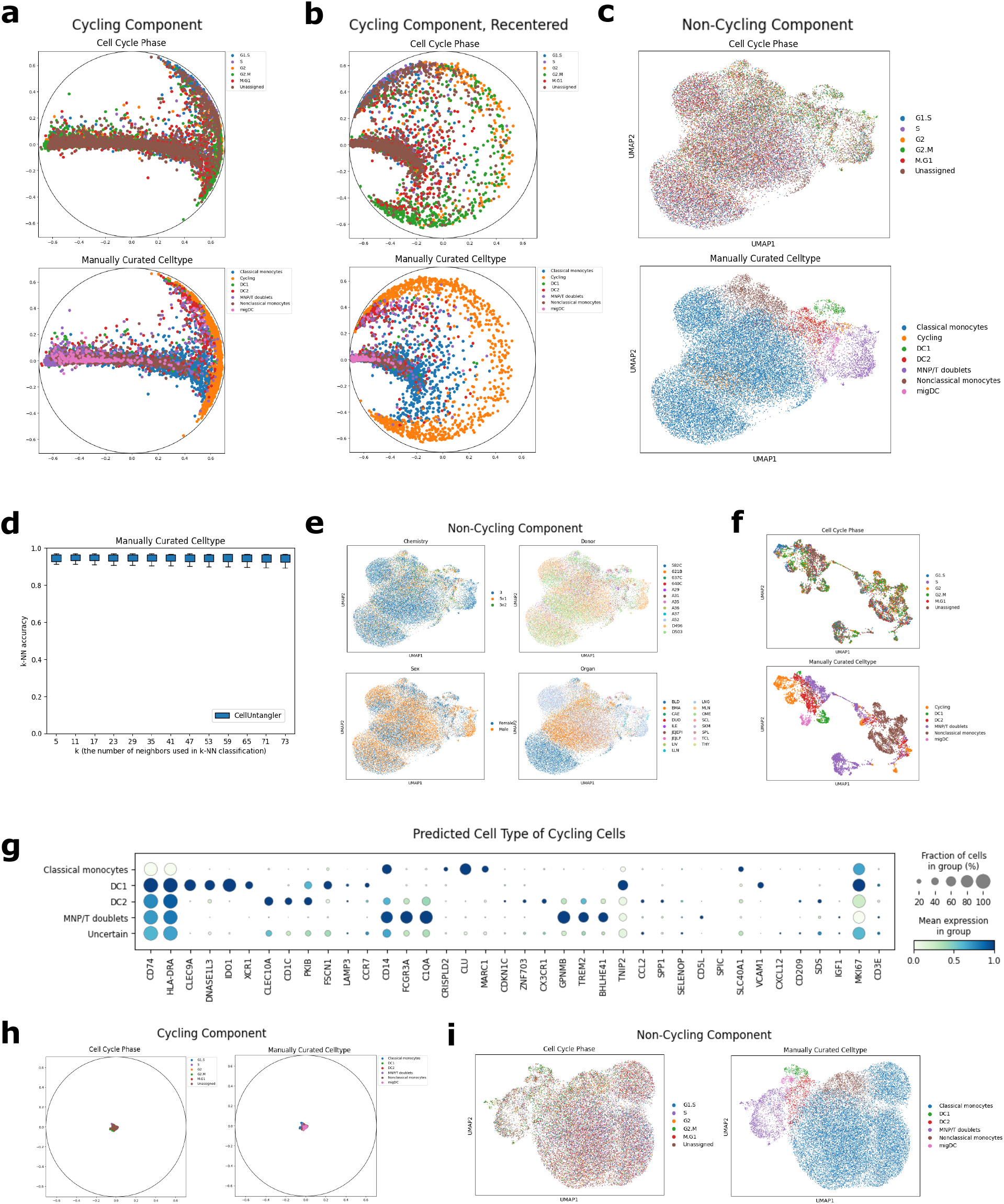
CellUntangler outputs embeddings that separately capture the cell cycle and cell type. Embeddings capturing cell type information no longer contain the effects of the cell cycle. (**a**) The embeddings of the first component, **z**^1^, before recentering, and (**b**) after recentering the origin to (−0.55, 0.0). (**c**) The embeddings of the second component, **z**^2^, after projecting to 2D using UMAP. (**d**) The *k*-NN accuracies on the manually curated cell type using **z**^2^, excluding cycling cells. Boxplots depict the medians and the interquartile ranges (IQRs). The whiskers are the lowest datum still within 1.5 IQR of the lower quartile and the highest datum still within 1.5 IQR of the upper quartile. (**e**) The embeddings of the second component, **z**^2^, colored by the batches present in the dataset. (**f**) UMAP of the cells after enhancing the data for the cell cycle signal using a cyclic topological prior with scPrisma. (**g**) Marker gene expression for predicted cell type of the cycling cells. (**h**) The embeddings of the first component, **z**^1^, and (**i**) the second component, **z**^2^, when the dataset is composed of only non-cycling cells.

Conversely, in the second component (**z**^2^) all cells, including cycling cells, were separated by cell type (**Fig. 4c**). Notably, cycling cells of all phases were integrated with other cell types (**Supplementary Fig. 3a, b**), especially monocytes and DC2s. The small patch of cycling cells likely represents macrophages (discussed below). To validate that the second component **z**^2^ successfully captured cell type information, we performed *k*-NN classification on the manually curated cell types (**Fig. 4d**), achieving a high accuracy of 0.94 for different *k*s, even though, as expected, these myeloid cells do not form very discrete clusters. (Cycling cells were excluded from the *k*-NN experiments.)

Moreover, the second component mitigated well technical batch effects, such as different sequencing techniques, while preserving biological variation, such as organ (**Fig. 4e**). In contrast, other methods, struggled with this use case. For example, scPrisma incorrectly placed non-cycling cells on the circular manifold when attempting to enhance the cell cycle signal (**Fig. 4f**); the standard Scanpy^34^ pipeline had cycling cells as distinct clusters (albeit in proximity to specific cell types) (**Supplementary Fig. 4a**) and strong batch effects remained (**Supplementary Fig. 4b**).

We used the *k*-NN classifier to determine the identity of the cycling cells (Methods), and confirmed those predictions by examining known marker gene expression (**Fig. 4g** and **Supplementary Fig. 5a**). Cycling cells predicted to be classical monocytes expressed *CD14* ; predicted DC1s expressed *XCR1* and *CLEC9A*; and predicted DC2s expressed *CD1C* and *CLEC10A*. We hypothesized that cycling cells predicted as MNP/T doublets or uncertain (when classifiers with different values of *k* did not predict the same cell type) were macrophages, because the original dataset contained macrophages which we filtered out prior to running CellUntangler. The macrophages in the original dataset included alveolar macrophages (from the lung), intermediate macrophages (lung), intestinal macrophages (jejunum), and erythrophagocytic macrophages (spleen, liver, mesenteric lymph nodes) (**Supplementary Fig. 6a**)^35^. These cells did not express the expected T-cell marker genes for MNP/T doublets. Most of these predicted MNP/T doublets were found in the lung and expressed markers of alveolar macrophages, including *GPNMB* and *TREM2* (**Supplementary Fig. 6b**). These cycling cells may have been classified as MNP/T doublets because some of the MNP/T doublets may contain macrophages, and the MNP/T doublets also expressed *GPNMB* and *TREM2* (**Supplementary Fig. 5a**). When examining the distribution of uncertain cells by organ (**Supplementary Fig. 6c**), we observed that cycling cells present in the jejunum expressed intestinal macrophage markers *CD209* and *IGF1*. Similarly, cycling cells present in the lung expressed *GPNMB* and *TREM2*, markers of alveolar macrophages.

Datasets may consist entirely of non-cycling cells. To evaluate this scenario, we ran CellUntangler on the myeloid cell dataset after excluding all cells annotated as cycling with the same settings. In **z**^1^, the cells no longer formed a circle, as the non-cycling cells were placed in the origin (**Fig. 4h**), whereas cell type information was still well captured in **z**^2^ (**Fig. 4i**).

### CellUntangler captures differentiation trajectories

Thus far, we have used the Euclidean space to capture the non-cycling component. However, some programs, such as those that govern cell differentiation trajectories, may be better represented in the hyperbolic space^22, 23^. To demonstrate the versatility of CellUntangler, we tested it on a dataset of mouse pancreatic cells^36^, which included both cycling and non-cycling cells, undergoing differentiation. This dataset exhibited two distinct signals: the cell cycle signal from the cycling cells and a strong differentiation trajectory signal from cycling to differentiated cells.

To capture these signals, we employed CellUntangler with two hyperbolic spaces: one for the cell cycle (**z** ∈ ℍ^2^, with the RoWN) and another for differentiation trajectories (**z** ∈ ℍ^2^). As expected, the first component (**z**^1^) successfully captured the cell cycle signal (**Fig. 5a**). After recentering, the cycling cells were arranged in a circular pattern around the non-cycling cells (**Fig. 5b**; Methods). Importantly, the second component (**z**^2^) effectively captured the differentiation trajectory. In the pancreas, ductal cells gradually induce Ngn3 first to become Ngn3 low cells followed by Ngn3 high cells and finally, the Ngn3 high cells express the transcription factor Fev (pre-endocrine cells) and differentiate into alpha, beta, delta, or epsilon cells (**Fig. 5c**).

**Figure 5:**
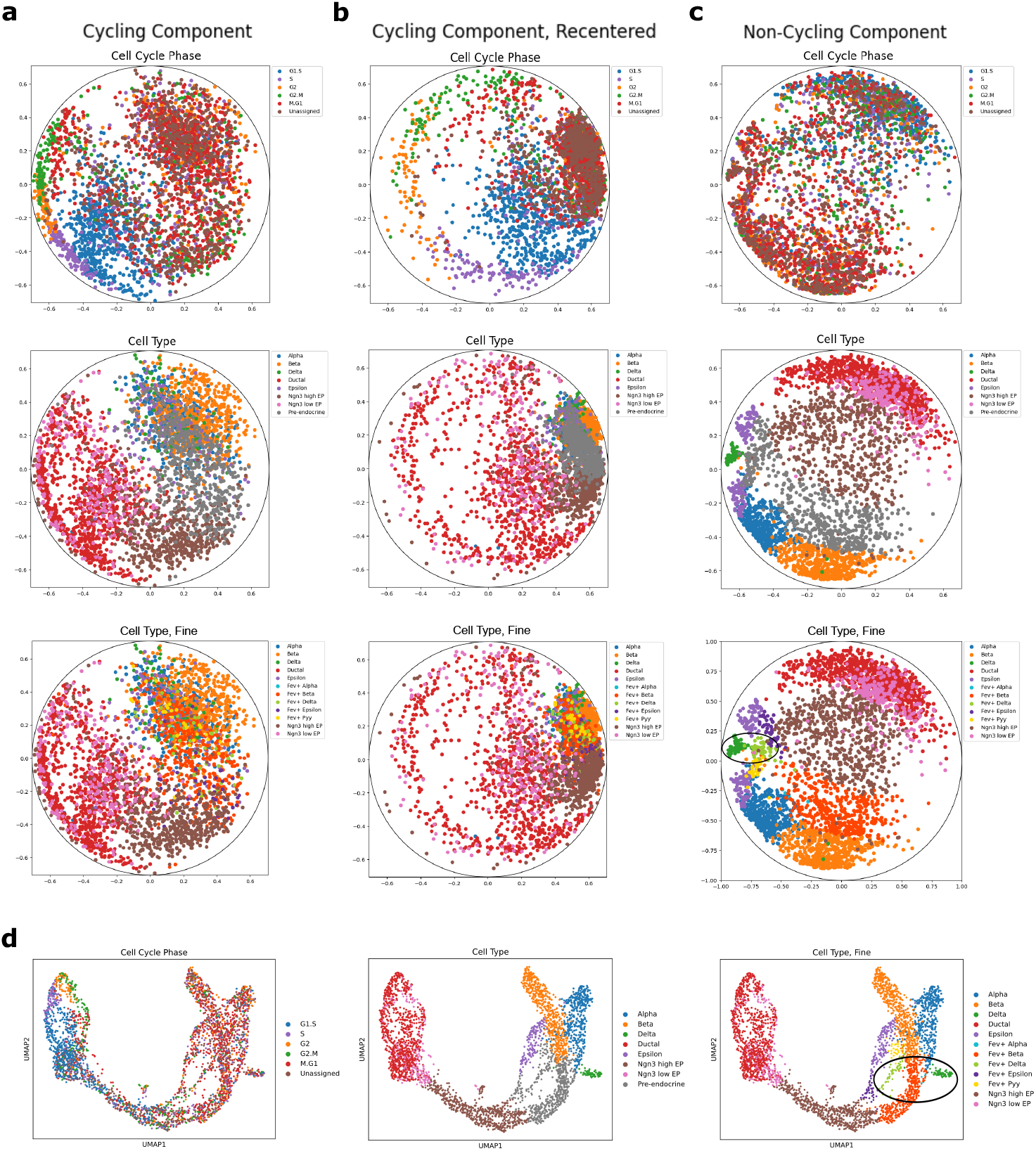
CellUntangler separates the cell cycle into one component and the differentiation trajectory into the other component. (**a**) The embeddings of the first component, **z**^1^, before recentering and (**b**) after recentering the origin to be (0.4, 0.1). (**c**) The differentiation trajectory is captured in the second component, **z**^2^. (**d**) A previously published UMAP^37^ of the same cells input to CellUntangler from the mouse pancreas dataset. In (**c**) and (**d**), the transition from Fev^+^ delta to mature delta cells is circled in black.

Further analysis of the pre-endocrine cells^36^ determined which cell type the cells were committed to. As expected, the Fev^+^ alpha cells were placed before the alpha cells and similarly, for the beta and delta cells. However, in the UMAP embeddings from studying pancreas cell development^37^, the differentiation signal is entangled with the cell cycle signal and notably, the fine-grained developmental trajectories are more clearly delineated in CellUntangler’s **z**^2^ embeddings compared to the UMAP embeddings (**Fig. 5d**). For example, the Fev^+^ delta cells were placed before the mature delta cells in the hyperbolic embeddings but separated by the Fev^+^ beta cells in the UMAP embeddings. Importantly, the epsilon cells were split into two groups: *Acsl1* ^+^ epsilon cells^38^ and *Acsl1* ^−^ epsilon cells (**Supplementary Fig. 7a**). The *Acsl1* ^+^ epsilon cells were placed following the Fev^+^ epsilon cells and the *Acsl1* ^−^ epsilon cells were placed following the Fev+ Pyy cells. Previous studies^36, 37, 39^ also place the Fev^+^ epsilon and Fev^+^ Pyy cells before the epsilon cells, but the pattern is much clearer in CellUntanger’s embeddings.

### CellUntangler captures and filters spatial zonation signals

While the cell cycle is an illustrative and common form of a predominant signal, CellUntangler can generalize to other cases where we wish to capture, distinguish, or remove different processes in single cells.

To demonstrate this broader applicability, we applied CellUntangler to a dataset of hepatocytes profiled from the mouse liver^40^. Hepatocytes in the liver have both zonation programs, reflecting their spatial location (and distinct functional requirements) in the liver lobules^41^, and circadian programs, reflecting timing in the light-dark cycle. In this dataset, cells from across the lobules were sampled at four evenly spaced time points throughout the day.

We used the Euclidean space to capture zonation in the first component, **z**^1^ ∈ 𝔼^2^, relying on 15 spatial marker genes^11,42^ and the second component, **z**^2^ ∈ 𝔼^10^, for the non-zonal (circadian) signal. As expected, in **z**^1^ cells were organized into different zonation layers, and mixed by time point (**Fig. 6a**), whereas in **z**^2^, the cells separated by time point only and not by layer (**Fig. 6b**).

**Figure 6:**
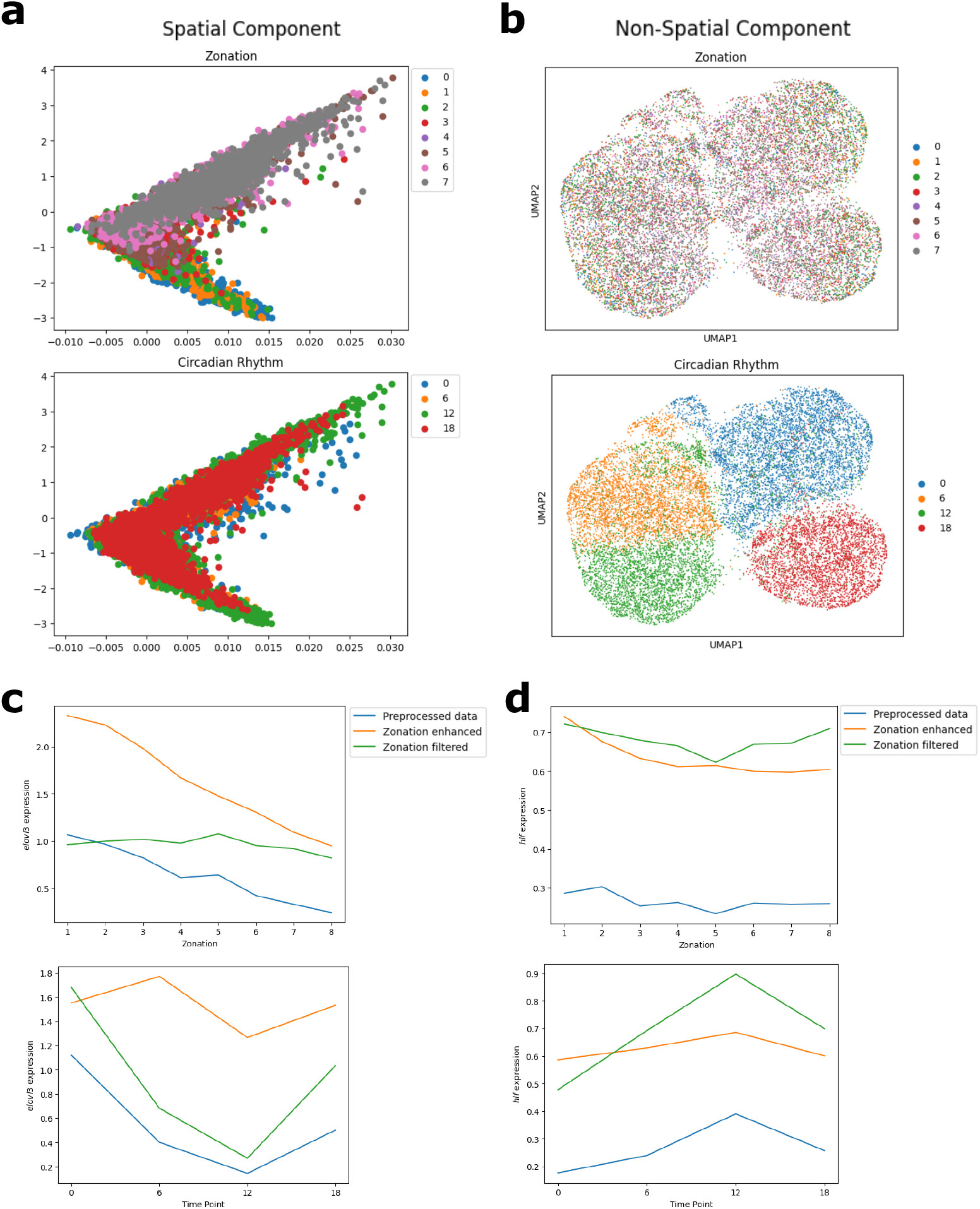
Spatial and temporal signals are captured and separated with CellUntangler. (**a**) The embeddings of the first component, **z**^1^; (**b**) the embeddings of the second component, **z**^2^, projected to 2D using UMAP. (**c**) Gene expression of *elovl3* across zonation (top) and circadian time point (bottom); (**d**) Gene expression of *hlf* across zonation (top) and circadian time point (bottom).

We examined the expression of spatiotemporal informative genes (*elovl3* and *hlf*) in the preprocessed data, the data enhanced for zonation (passing **z**^1^ to the decoder), and the data with the zonation signal filtered (passing **z**^2^ to the decoder). For *elovl3*, differential expression across zonation layers (**Fig. 6c**, top) was retained for the zonation-enhanced data and flattened for the zonation-filtered data. Across circadian rhythm time points (**Fig. 6c**, bottom), differential expression of *elovl3* was flattened for the zonation-enhanced data and retained for the zonation-filtered data. Gene expression varied similarly for *hlf* across zonation layers (**Fig. 6d**, top) and circadian rhythm time points (**Fig. 6d**, bottom).

## Discussion

We introduced CellUntangler, a deep generative model designed to capture and separate biological signals in cells by embedding them into a decomposed latent space, where each subspace can be Euclidean, hyperspherical, or hyperbolic. CellUntangler allows us to obtain separate embeddings for each signal, and we demonstrated its effectiveness on datasets consisting of the cell cycle signal along with genetic or cell type differences or differentiation trajectories, and with both zonation and circadian signals.

CellUntangler works well on datasets composed of only cycling cells. On the HeLa cell dataset, it successfully separated cells by cell cycle stage and by genotype, and when comparing *k*-NN accuracies, it showed favorable performance for smaller values of *k* and comparable results for larger values. Although Revelio performed similarly well, it required more cycling genes and prior knowledge of the cell cycle stage for each marker gene. CellUntangler only requires knowledge of the marker genes for the signal. scPrisma, which does not require any prior knowledge of marker genes, demands more hyperparameter tuning for the test dataset. On the dataset of mouse embryonic stem cells, the cells proceeded in the order of pseudotime, while no such ordering was observed in the second latent subspace. Visualizing the data in the Poincaré disk also provides a convenient measure of pseudotime, based on the angle from the origin. We found that our pseudotime estimation correlated well with the ground truth, demonstrating the effectiveness of the hyperbolic space with the RoWN in capturing the cell cycle signal. A key strength of CellUntangler is its ability to handle both cycling and non-cycling cells within the same dataset. Other methods may struggle in this regard, as they typically assume all cells follow the cell cycle trajectory. CellUntangler, however, distinguishes cycling cells from non-cycling cells by positioning cycling cells along the edge of the Poincaré disk.

Importantly, CellUntangler captures multiple signals in simultaneous representations that capture and filter out the desired signals, reducing the number of steps required, and the second component can retain relevant cell information while filtering out the prominent effects of another process, such as the cell cycle or zonation. In the immune cell dataset, cell type information is preserved, allowing for accurate identification of the cell types of cycling cells, which can help interpret scRNA-seq datasets. In the pancreatic cell dataset, it captures a differentiation trajectory. In the myeloid cell atlas, it handled batch effects, ensuring that irrelevant information is not captured in the latent spaces, eliminating the need for a separate step to remove such noise.

Crucially, CellUntangler can distinguish, capture, and filter other signals, such as zonation and circadian rhythms in the mouse liver cell dataset, and only knowledge of some marker genes for the primary signal is required. Additionally, CellUntangler can be extended to use more complex latent spaces with multiple subspaces. For example, the cell cycle could be represented by embedding cells along a cylinder, the product space of a circle and a line. Analyses with markers for organ-specific features and multiple subspaces could also be used to tackle variation from the organ in the immune cells dataset. We anticipate that CellUntangler will be a powerful tool for downstream analysis and filtering of various signals, and its flexibility in latent space will enable us to gain deep insights into single-cell data.

## Methods

### The CellUntangler model

CellUntangler isolates and reveals biological processes that might obscure one another. It takes as input a gene expression count matrix and a set of marker genes for a known signal. It separates the known signal from other signals present in the dataset, by embedding each cell into a latent space consisting of several subspaces, with each subspace having the appropriate geometry to capture the signal of interest.

A scRNA-seq dataset 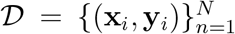, where *N* is the number of cells, **x**_*i*_ ∈ ℝ^*G*^ is a Unique Molecular Identifier (UMI) count vector, *G* is the number of genes, and **y**_*i*_ is a categorical vector of the batch or batches associated with **x**_*i*_, is provided as input to CellUntangler. **x**_*i*_ is log-transformed prior to input into the model. Single-cell data intrinsically has low-dimensionality^22^, so we assume that the distribution of **x**_*i*_ is governed by a lower-dimensional vector **z**_*i*_. Accordingly, the joint distribution can be factorized as follows

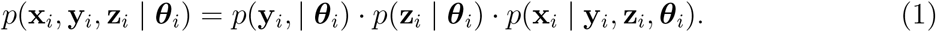

Here, *p*(**y**_*i*_ | ***θ***_*i*_) is the categorical distribution, and *p*(**z**_*i*_ | ***θ***_*i*_) is the prior distribution for **z**_*i*_. The parameter set ***θ***_*i*_ represents the parameters of each distribution. Since the distributions are different, ***θ***_*i*_ will differ for each distribution. For example, if *p*(**z**_*i*_ | ***θ***_*i*_) follows a multivariate normal distribution, ***θ***_*i*_ would be ***µ***_*i*_ and **Σ**_*i*_. To capture the complex nonlinear relationship between **z**_*i*_ and **x**_*i*_, CellUntangler uses neural networks to parametrize the distribution *p*(**x**_*i*_ | **y**_*i*_, **z**_*i*_, ***θ***_*i*_). The use of a neural network for *p*(**x**_*i*_ | **y**_*i*_, **z**_*i*_, ***θ***_*i*_) makes the computation of the posterior *p*(**z**_*i*_ | **x**_*i*_, ***θ***_*i*_) intractable, so variational inference is used with the variational distribution *q*(**z**_*i*_ | **x**_*i*_, ***ϕ***_*i*_), an approximation of the true posterior where the variational parameters ***ϕ***_*i*_ are output by a neural network.

CellUntangler consists of two neural networks: the inference network (encoder) and the model network (decoder), forming a variational autoencoder (VAE)^12^ architecture. In a conventional VAE, the latent space is typically a single space with a constant curvature, such as the Euclidean, hyperspherical^20^, or hyperbolic latent spaces^21^. Different dimensions of the latent space are treated equally, and they jointly learn representations of the input data, without explicitly encoding distinct signals into different dimensions of the latent space. In contrast, CellUntangler allows for a product of spaces, where each subspace can have a distinct curvature, either Euclidean, hyperspherical, or hyperbolic with varying dimensionality for *k* subspaces, as in the mixed-curvature variational autoencoder (MVAE)^43^ framework. This design enables CellUntanger to potentially capture and disentangle distinct signals. To help capture biological processes of interest, e.g., cell cycle, stress, and cell type information, we decompose the latent vector 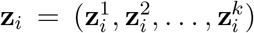, the concatenation of vectors 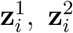, and 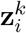, such that each compartment 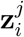 presents a biological process, e.g., cell cycle. The posterior can now be written as 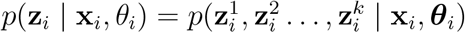. The choice of geometry for each latent subspace is tailored to the specific signal being modeled. For example, a hyperbolic space might be selected to capture differentiation trajectories, which often exhibit tree-like structures.

To untangle the confounding signal from other signals in the dataset, known marker genes for that signal are required as input. For many biological processes of interest, this prior knowledge (e.g., stress, interferon response)^44, 45^ is available or can be a set of marker genes that can be defined by differential expression analysis. For instance, cell cycle marker genes can be obtained from databases, by literature curation (manually or with a large language model), or as genes upregulated in cycling vs. non-cycling cells (data-driven). For simplicity of discussion, **z**_*i*_ is decomposed into two latent components. However, the **z**_*i*_ in CellUntangler can be decomposed into multiple components (**Supplementary Note 1**).

To use 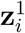 to capture the known signal (e.g., cell cycle) and 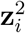 to capture the other signals (e.g., cell type) that may be confounded by the known signal, **x**_*i*_ is decomposed into two UMI count vectors, one for the marker genes, 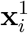, and one for the non-marker genes, 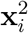. From 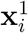 and 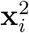, the encoder outputs the parameters of 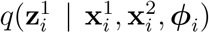 and 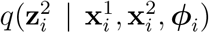. When computing 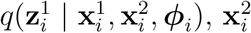 is masked (set to zero), while when computing 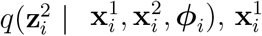 is masked.

*x*_*ij*_, the UMI count of gene *j* in cell *i*, is assumed to follow a negative-binomial distribution, based on previous work^46–48^. When decomposing 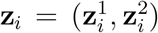 and assuming there are *m* marker genes associated with the signal encoded in 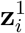,

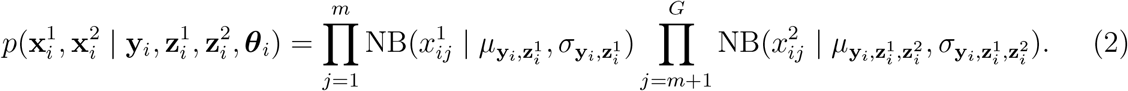

Only 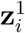 is used to reconstruct 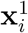, while both 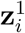 and 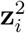 are used to reconstruct 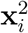, as 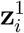 may be informative of 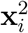.

Independence between the latent representations would allow better separation of biological signals into 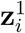 and 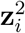. However, backpropagation may cause information from 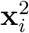 to be captured in 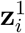, as 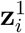 and 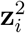 are not conditionally independent given 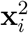. As the same gene may have activities related to more than one process, this matters particularly when using a larger set of marker genes, since some may be less specific to the desired signal. To obtain more independent representations, when the decoder is trained, 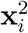 is reconstructed in two ways. The first uses 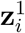 and 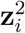 to reconstruct 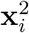 (**Fig. 7a**). The second stops the gradient for 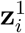 so that it is fixed before reconstructing 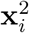 (**Fig. 7b**). The number of epochs for training with stop gradient for 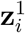 in reconstructing 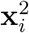 is a hyperparameter in the model.

**Figure 7:**
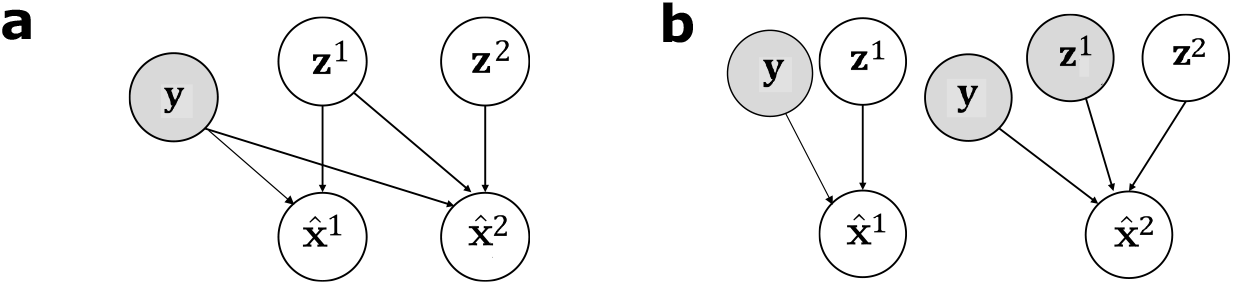
Probabilistic graphical models for CellUntangler. The probabilistic graphical model for CellUntangler when the latent representation is decomposed into **z**^1^ and **z**^2^, (**a**) when stop gradient is not used, and (**b**) when it is used.

To find the parameters for the inference neural network and model network that maximize the objective function, the evidence lower bounds (ELBO) is used,

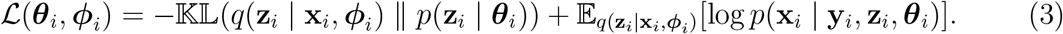

The ELBO requires sampling from the variational distribution and evaluating the density of samples from distributions of the Euclidean, hyperspherical, and hyperbolic spaces. The Euclidean space is defined as 𝔼^*d*^ = ℝ^*d*^ for *K* = 0 where *d* is the dimension and *K* is the curvature. The hyperspherical space has constant positive curvature and is defined as

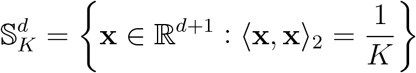

for *K >* 0, where 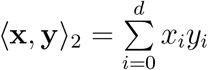 is the standard inner product. The hyperbolic space has constant negative curvature and is defined as

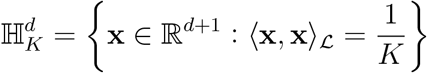

for *K <* 0 where 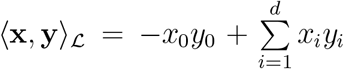 is the Lorentz inner product, and the first elements *x*_0_ *>* 0 and *y*_0_ *>* 0.

For each of the three constant curvature spaces, a tangent vector space is defined at every point of the space, so we use wrapping to obtain the wrapped normal distribution^21^ for the prior distribution *p*(**z**_*i*_ | ***θ***_*i*_) and the variational distribution *q*(**z**_*i*_ | **x**_*i*_, ***ϕ***_*i*_). To sample from the wrapped normal distribution with mean ***µ*** and covariance matrix **Σ**, denoted by 𝒲𝒩(***µ*, Σ**), the origin ***µ***_0_ for each space is defined as: For the Euclidean space, ***µ***_0_ = **0** ∈ ℝ^*d*^. For the hyperspherical and hyperbolic space, 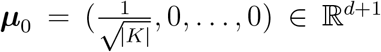. Next, 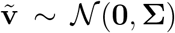 where 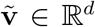 is sampled, and, for the hyperspherical and hyperbolic space, concatenated so that the first element is 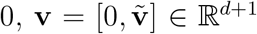. From the definition of the origin for each space, ⟨***µ***_0_, **v**⟩_2_ = 0 and ⟨***µ***_0_, **v**⟩_ℒ_ = 0, therefore 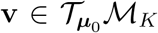 where 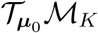 is the tangent space at ***µ***_0_ of the manifold with curvature *K*. Parallel transport is used to transport **v** to the tangent space at 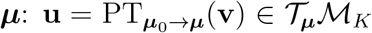. Finally, the exponential map is used to map **u** from the tangent space to the manifold: 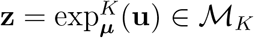.

To calculate the probability density of the samples, the reverse procedure is used. Given a sample **z**, the logarithmic map is used to map the vector from the manifold to the tangent space at 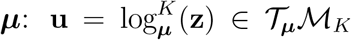. Parallel transport is used to transport **u** from the tangent space at ***µ*** to the tangent space at 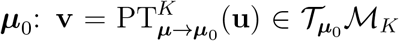. The density is computed as 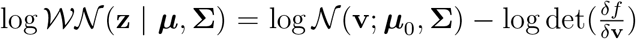, where 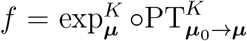. The KL divergence can be estimated using Monte Carlo integration.

### The rotated hyperbolic wrapped normal distribution

When using the hyperbolic space to capture the cell cycle signal, the rotated hyperbolic wrapped normal distribution (RoWN) is used as the variational distribution for the latent cell cycle signal of each cell. To sample from 𝒲𝒩(***µ*, Σ**), sampling must first be done from 𝒩 (**0, Σ**), typically with a diagonal covariance matrix **Σ** = diag (***σ***). As the covariance matrix is diagonal, the covariance structure will have principal axes parallel to the standard bases. The principal axes will also go through the origin since sampling is from a normal distribution with a mean of **0**. As a result, the principal axes can be viewed as straight lines that pass through the origin, *l*_**s**_(*t*) = *t***s** ∈ ℝ^*d*^, where **s** ∈ ℝ^*d*^ is a directional vector. Straight lines that pass through the origin become geodesics in the hyperbolic space (using the exponential map) and remain geodesics when projected to a Poincaré ball^27^. Thus, the principal axes for the diagonal covariance matrix, for the normal distribution with a mean of **0**, in the Euclidean space become geodesics in the hyperbolic space. The vector **s** will become a tangent vector of the geodesic resulting from projecting the principal axis to the Poincarè ball on the projected point ***µ***. As **s** is parallel to the standard bases and a tangent vector of the projected principal axes, the projected principal axes in the hyperbolic space are also locally parallel to the standard bases. The principal axes determine the covariance structure, so local variation can only be modelled parallel to the standard bases, instead of pointing in the radial direction. This is problematic because cycling cells should form a circle of approximately equal distance to the origin in Poincaré disk, such that their angular distance reflects their cell cycle phase, and two cells of the same cell cycle phase should have approximately similar angles. Variations in the radial direction of the Poincaré disk should model cell cycle-independent factors, such as noise. In contrast, the hyperbolic space with the RoWN rotates the principal axes of the covariance matrix so that local variation can be modeled in the radial direction.

Given ***µ*** ∈ ℍ^*d*^ so ***µ*** ∈ ℝ^*d*+1^, two vectors are obtained: **x** = [±1, …, 0] ∈ ℝ^*d*^, where ± is determined by the sign of the first element of ***µ*** and 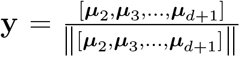, where *µ*_*i*_ is the *i*th element of ***µ***. Given the two row vectors, **x** and **y**, the rotation matrix is computed as:

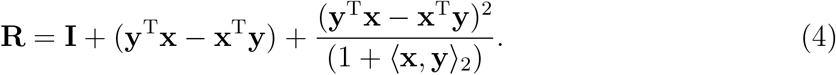

⟨**x, y**⟩_2_ is the standard inner product. Next, **Σ** is rotated to obtain 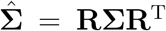. The same steps as before are followed but with sampling from 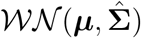 instead of 𝒲𝒩 (***µ*, Σ**), allowing sampling to remain efficient and probability density evaluation to remain tractable.

### Recentering the origin

The origin is recentered by the Mobius addition, which for two vectors 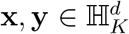 with curvature *K* is defined as:

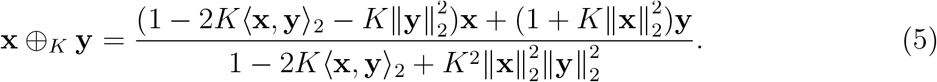

When embedding datasets containing cycling and non-cycling cells, cycling cells are embedded along the edge of Poincaré disk, while non-cycling cells are placed through the rest of the disk. After locating cycling cells along the edge of the disk, the leftmost cycling cell is estimated to have coordinates [*x*_*l*_, *y*_*l*_]^T^, and the rightmost cycling cell to have coordinates [*x*_*r*_, *y*_*r*_]^T^. The estimated topmost cycling cell has coordinates [*x*_*t*_, *y*_*t*_]^T^, and the bottom-most cycling cell has coordinates [*x*_*b*_, *y*_*b*_]^T^. The center of cycling cells can then be estimated as 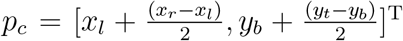. We recommend choosing the new origin, *o*_*n*_, as the point diagonally across from *p*_*c*_ near the edge of the Poincaré disk, so that when a line is drawn from *p*_*c*_ to *o*_*n*_, approximately half of the cycling cells line above and below the line. The line should be approximately perpendicular to the curve of the cycling cells.

### Model structure

Due to the sparsity of single-cell data, the softmax activation function is used on the outputs of the decoder to output a vector of positive numbers that sum to one and encourage sparse outputs. The softmax outputs are multiplied by the sum of UMI counts for each cell to obtain the final means of the negative binomial distributions for that cell. The Gaussian Error Linear Unit (GELU)^49^ is used as the activation function for the hidden layers.

The encoder consists of three layers (128-64-32) and the decoder consists of two layers (64-128). The weights of the layer that outputs the parameters of the cell cycle component are set with Xavier normal initialization^50^, while the remaining weights are set with the default initialization. A curvature of -2 is used for the cell cycle component. The dimensionality of the cell cycle component is set to two to facilitate visualization and to embed the cycling cells in a circular structure. The model is trained for 500 epochs using mini-batches of size 128 with the AdamW^51^ optimization algorithm, a learning rate of 0.001 and a weight decay of 0.01.

The number of epochs during which the gradient is stopped for 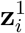 in the reconstruction of 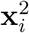 is a hyperparameter. For the mouse pancreas dataset, it was stopped for all 500 epochs due to the presence of a strong differentiation signal. For all other datasets, the gradient for 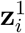 was not stopped at any point.

### Cell cycle genes

An input list of 229 human cell cycle genes was derived from a single-cell esophageal mucosal cell atlas^52^ as follows. For 10 of 11 cycling cell subsets, including cycling fibroblasts, endothelial cells, pericytes, mast cells, macrophages, dendritic cells, B cells, CD4^+^ T cells, CD8^+^, and NK cells, differential expression analysis was performed to find genes differentially expressed between cycling cells and their non-cycling counterparts in the same cell type, *e*.*g*., cycling B cells compared to non-cycling B cells. Cycling signature genes were defined as genes that were upregulated (Bonferroni adjusted *p*-values *<* 0.01, log2 fold-change *>* 0.5, expressed in at most 15% of the non-cycling cell subsets) in at least five of the cycling subsets, to a total of 194 genes. These 194 cycling signature genes were combined with 97 cell cycle marker genes from an earlier study^53^ (62 overlapping genes), resulting in a final list of 229 cell cycle signature genes. For differential expression analysis, we used the Mann-Whitney *U* test from Seurat.

For mouse datasets, orthologs of the human cell cycle genes were identified. Orthologs were obtained from the Mouse Genome Informatics (MGI) Resource^54^. Of the 229 cell cycle signature genes, 16 were not present in the table used. Orthologs were found for 15 of the absent genes by using the search bar of the Mouse Genome Informatics (MGI) Resource. No orthologs were found for *FAM111B*. This resulted in a total of 256 mouse cell cycle signature genes as some human genes had more than one mouse ortholog.

When selecting highly variable genes in pre-processing, any cell cycle genes that are initially filtered out are added.

### Cell cycle pseudotime

A cell’s pseudotime ranges from [0^°^, 360^°^] or [0, 2*π*]. Pseudotime was obtained by first projecting 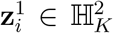 from Lorentz to Poincaré coordinates using the equation 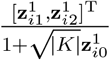 The pseudotime was measured as the angle of the projected Poincaré coordinates for each cell, relative to the origin of the Poincaré disk, (0, 0).

### Cell cycle reconstruction

To assess the quality of the cell cycle reconstruction, Pearson’s correlation coefficient of the pseudotime obtained from CellUntangler and the ground truth pseudotime was calculated. For the HeLa cell dataset, cell cycle time was used as the ground truth pseudotime. As the ground truth pseudotime and computed pseudotime may not be aligned, a rotation was applied, and, if necessary, the direction of the estimated pseudotime was flipped. The pseudotime with the highest Pearson correlation coefficient with the ground truth pseudotime was used to measure CellUntangler’s performance and for comparison of different methods. As the cell cycle angle was available for the HeLa cell dataset, the correlation between the sin of the ground truth angle and the sin of the measured pseudotime was also calculated. This additional correlation was used because cell cycle pseudotime is circular, meaning that values near 0 and 2*π* should be considered similar. Using the Pearson correlation coefficient directly on these values could result in a lower correlation. The use of the sin function helps to address this issue by mapping 0 and 2*π* to the same value, thereby improving correlation measurement.

### Cell identity of cycling cells

After separating the non-cycling cells from the cycling cells based off of the manually curated cell type^35^, *k*-NN was used to determine the identity of the cycling cells. The identity of a cell was labeled as uncertain if any one of the classifiers for different values of *k* did not predict the same cell type. We used values of 5, 11, 17, 23, 29, 35, 41, 47, 53, 59, 65, 71, and 73 for *k*.

### Batch effect removal

To address batch effects in the datasets, CellUntangler takes as input, the batch or batches. We did not use any batches for the HeLa dataset, mouse embryonic stem cells dataset, and mouse pancreas dataset. For the immune cells dataset, we used chemistry (Chemistry) and donor (Donor) as the batches. We used replicate (rep) as the batch for the mouse liver dataset.

### Parameter setting for other methods

To obtain the cell cycle stages for cells of a dataset, Revelio^28^ was run with the 800 genes included in the Revelio package, categorized into five groups: G1, G1.S, G2, G2.M, M.G1. Seurat^29–32^ for cell-cycle analysis only requires S and G2.M phase genes, and Seurat was run using all S and G2.M genes present in the cell cycle gene list we used for CellUntangler. Genes were assigned S or G2.M based on the gene list from Revelio. Similarly, Scanpy^34^ was run using all S and G2.M genes present in the cell cycle gene list we used for CellUntangler. SiFT was run by first normalizing the counts to 10,000 and then log-transforming the normalized counts with the 229 gene list defined above and a *k*-NN kernel based on the SiFT tutorial. scPrisma was run using the default parameters in their de novo reconstruction tutorial for cyclic signals. Classical monocytes were filtered out due to memory constraints in running scPrisma.

### Datasets

The HeLa dataset consists of 1,477 human cells profiled by Drop-seq^55^ and 4,545 genes were used based on the Revelio pipeline.

The mouse embryonic stem cells dataset consists of 5,637 cells and 11,625 genes profiled by 10x Chromium. The cells were filtered for quality by Riba *et al*.^6^. Genes were those present in both the raw and author-processed data^6^.

Human myeloid cells were profiled using a Chromium chip from 10x Genomics, the Single Cell 5’ Reagent (v1 and v2), and the 3’ Reagent (v3) Kit from twelve different donors. The dataset consists of 51,552 cells, and 29,376 cells of them were included in the analysis, selected as dendritic cells, monocytes, cycling cells, and MNP/T doublets. Highly variable genes were selected with Scanpy and any cycling genes not included in the highly variable gene list were added.

Mouse pancreatic stem cells were profiled using the 10x Chromium platform. The original dataset consists of 36,361 cells across four different embryonic stages, and the 3,559 cells from the E15.5 embryos selected in Zheng *et al*.^5^ were used, with the highly variable genes in the original study^36^, along with any cell cycle genes not included.

The mouse liver dataset was profiled using the 10x Chromium platform. 15,573 cells filtered for quality^11^ were used and 5,058 genes consisting of those remaining after filtering based on the consistency of the replicates^40^, along with any other zonation and circadian genes.

## Data Availability

All scRNA-seq datasets used in this study are publicly available. The HeLa cell dataset was acquired from the Gene Expression Omnibus (GEO) database under accession code GSE142277. The mouse embryonic cells dataset was acquired from GEO under accession code GSE167609. The immune cells dataset was downloaded from cross-tissue immune altas^35, 56^. The mouse pancreas dataset was acquired from GEO under accession code GSE132188. The mouse liver dataset was acquired from GEO under accession code GSE145197.

## Code Availability

The code is available at https://github.com/Ding-Group/CellUntangler.

## Acknowledgements

This work was supported by a Discovery grant from the Natural Sciences and Engineering Research Council (NSERC) of Canada, and a department startup fund from the University of British Columbia (to J.D.) as well as the Klaman Cell Observatory (A.R.). J.D. is a Canada Research Chair and is supported by the Canadian Institutes of Health Research through the Canada Research Chair Program. The computational resource is partially supported by the Canada Foundation for Innovation & John. R. Evans Leader Fund (to J.D.). This research was supported in part through the computational resources and services provided by Advanced Research Computing at the University of British Columbia.

## Author’s contributions

J.D. and A.R. conceived the study and developed the prototype model. J.D. and S.C. developed the model. S.C. conducted experimental analyses and interpreted the results with supervision from J.D. and A.C.. S.C., J.D., A.R., and A.C. wrote the manuscript.

## Supplementary Figures

**Supplementary Fig. 1:**
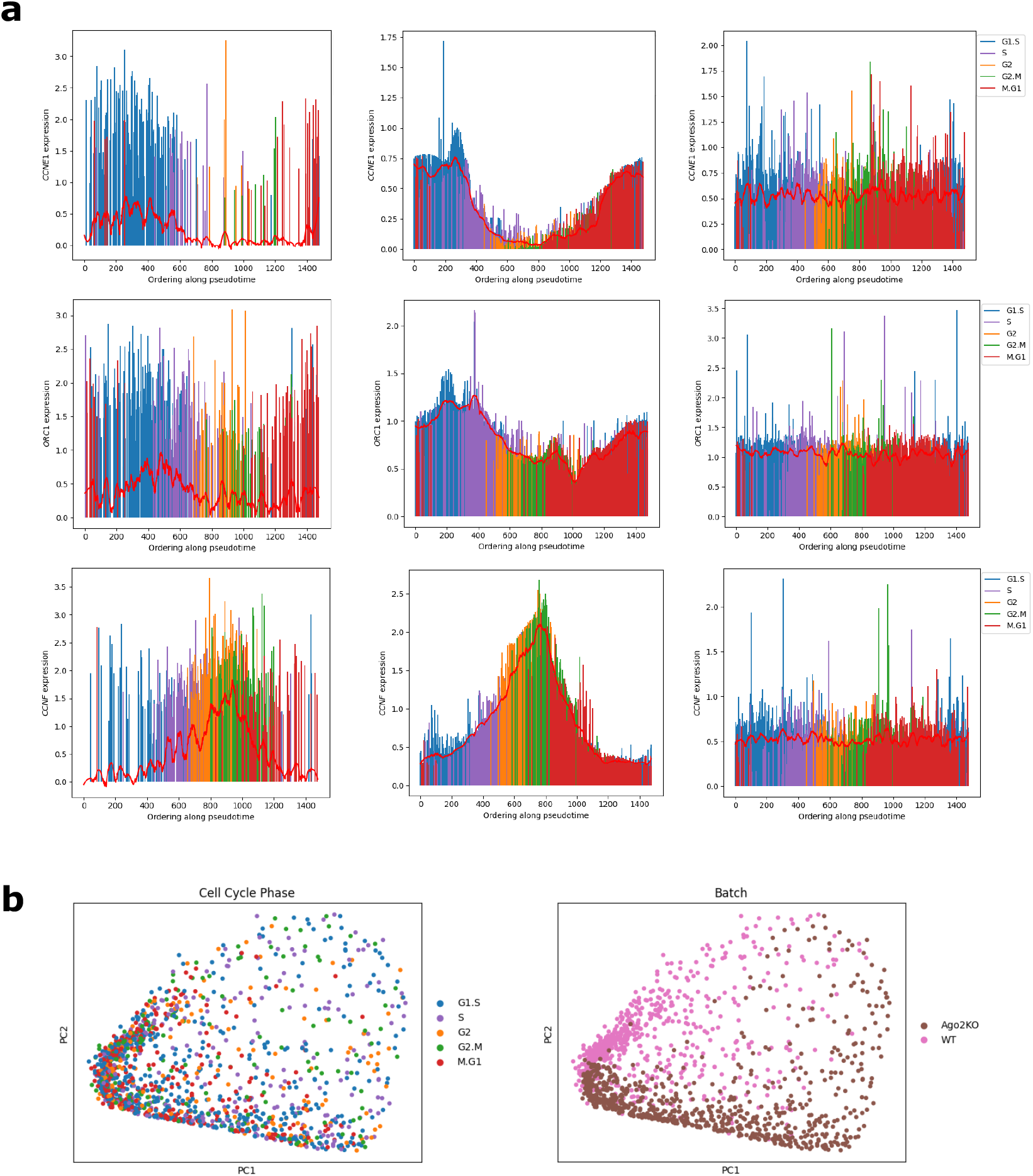
Analysis of the gene expression reconstructions obtained using z^1^ and z^2^ suggests capturing and removal of the cell cycle. (**a**) Gene expression of *CCNE1* (G1.S), *ORC1* (G1.S), and *CCNF* (G2) as a function of pseudotime obtained with the embeddings from CellUntangler using the original preprocessed data (size-factor normalized and log-transformed, left) and the reconstructed gene expression output from CellUntangler after decoding **z**^1^ (middle) and **z**^2^ (right). Cell cycle stages (G1.S, S, G2, M) are in parentheses. (**b**) PCA representation of the reconstructed data obtained by running **z**^2^ through CellUntangler’s decoder.

**Supplementary Fig. 2:**
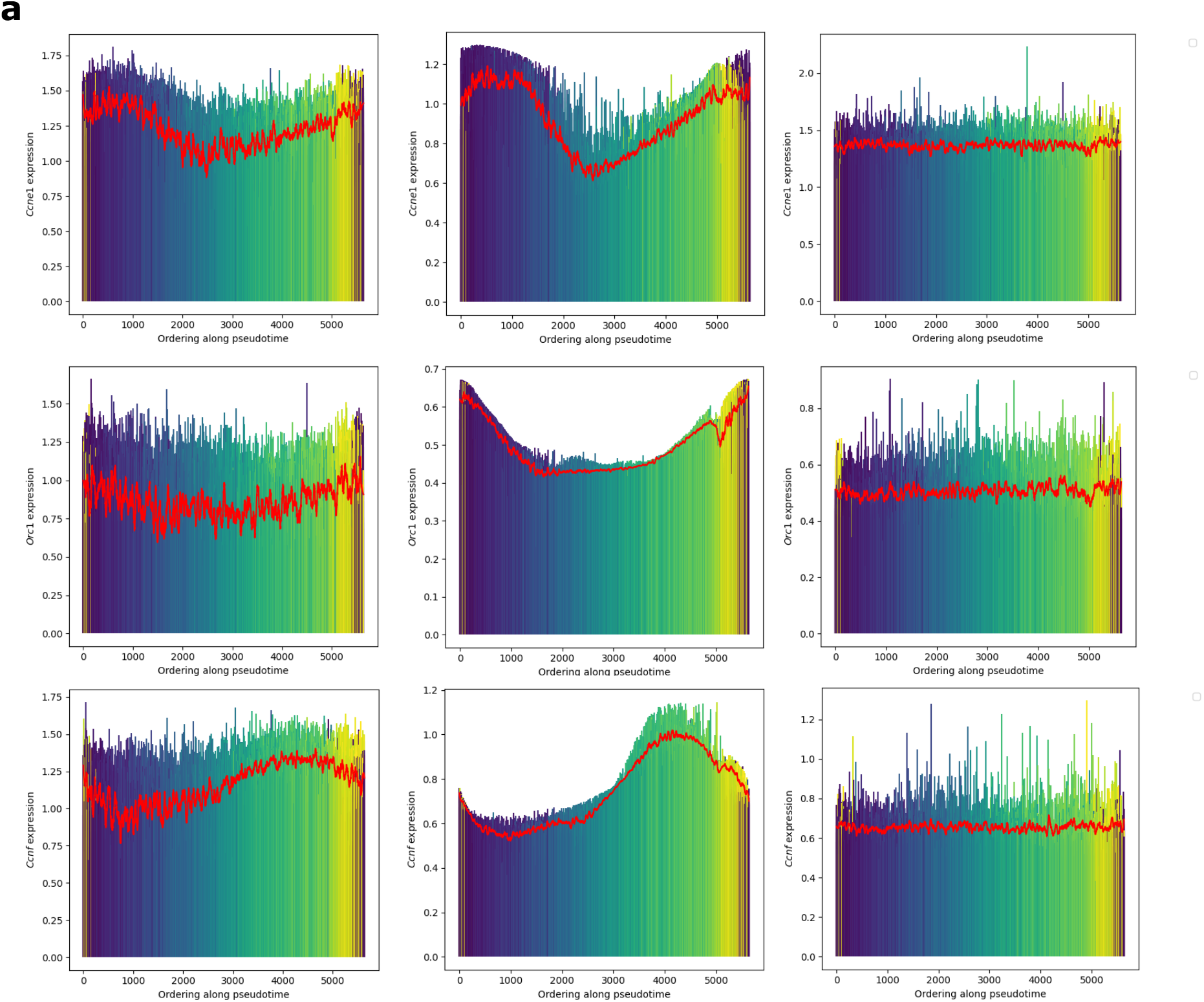
The effects of enhancing and filtering the cell cycle signal on putative cell cycle genes. (**a**) Gene expression of *Ccne1* (G1.S), *Orc1* (G1.S), and *Ccnf* (G2) as a function of pseudotime obtained with the embeddings from CellUntangler using the original preprocessed data (size-factor normalized and log-transformed, left) and the reconstructed gene expression output from CellUntangler after decoding **z**^1^ (middle) and **z**^2^ (right).

**Supplementary Fig. 3:**
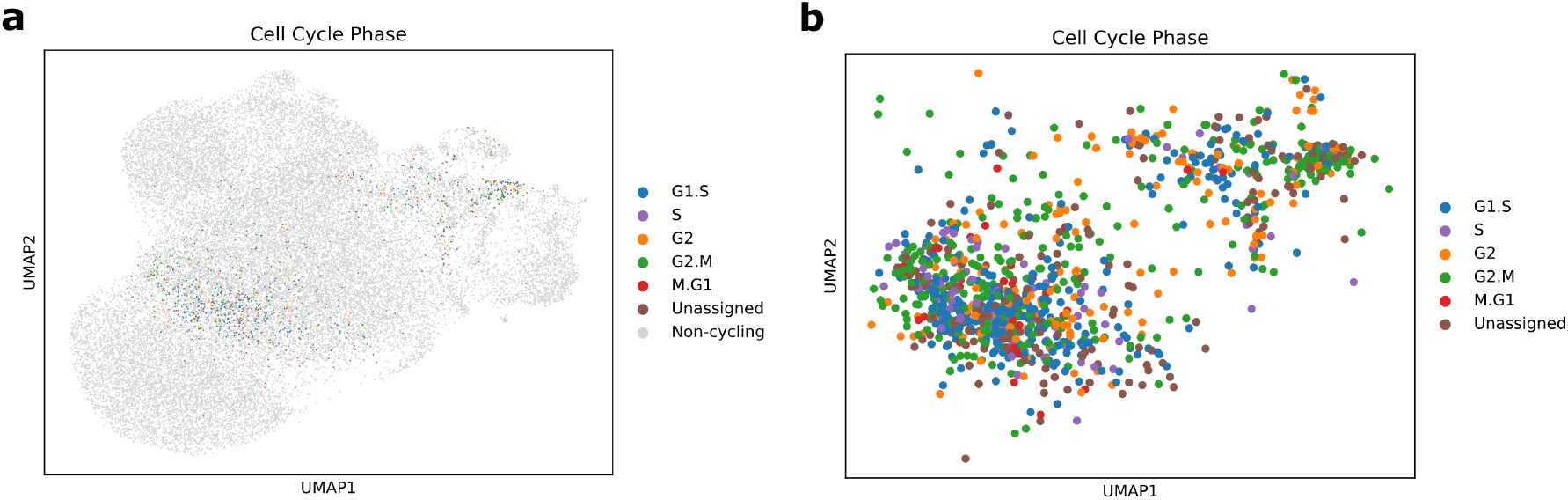
Visualization of CellUntangler’s second component demonstrated that cycling cells are well integrated by cell cycle phase. (**a**) The second component, **z**^2^, projected to 2D using UMAP when only cycling cells are colored by cell cycle phase while non-cycling cells are in gray. (**b**) The same UMAP as (**a**) but only the cycling cells.

**Supplementary Fig. 4:**
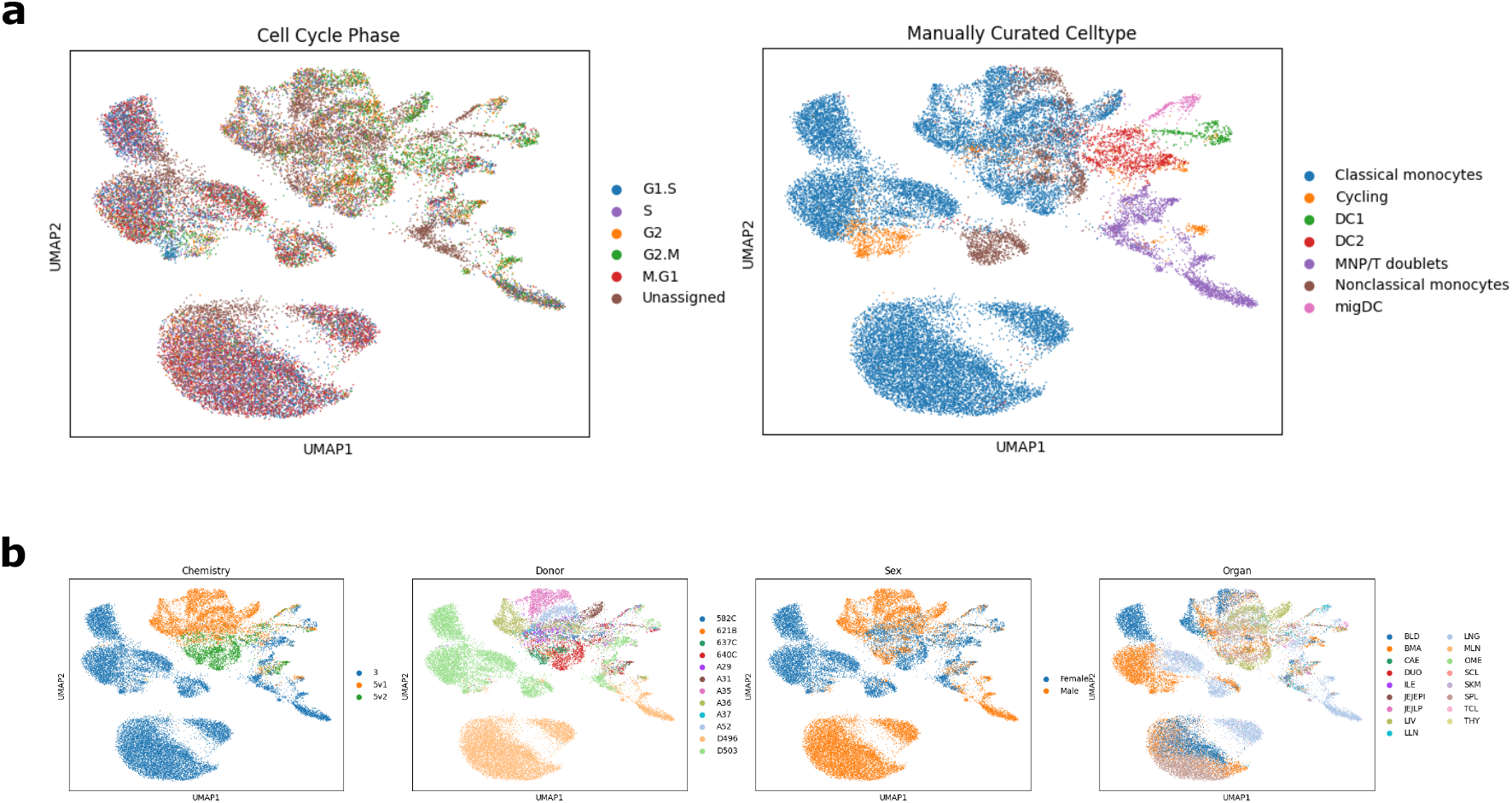
Applying Scanpy on the immune cells dataset to regress out the effects of the cell cycle. (**a**) UMAP visualizations colored by cell cycle phase (left) and manually curated cell type (right) after regressing out the cell cycle using Scanpy and performing PCA. (**b**) UMAP visualizations colored by different batches present in the dataset, after regressing out the cell cycle using Scanpy and performing PCA.

**Supplementary Fig. 5:**
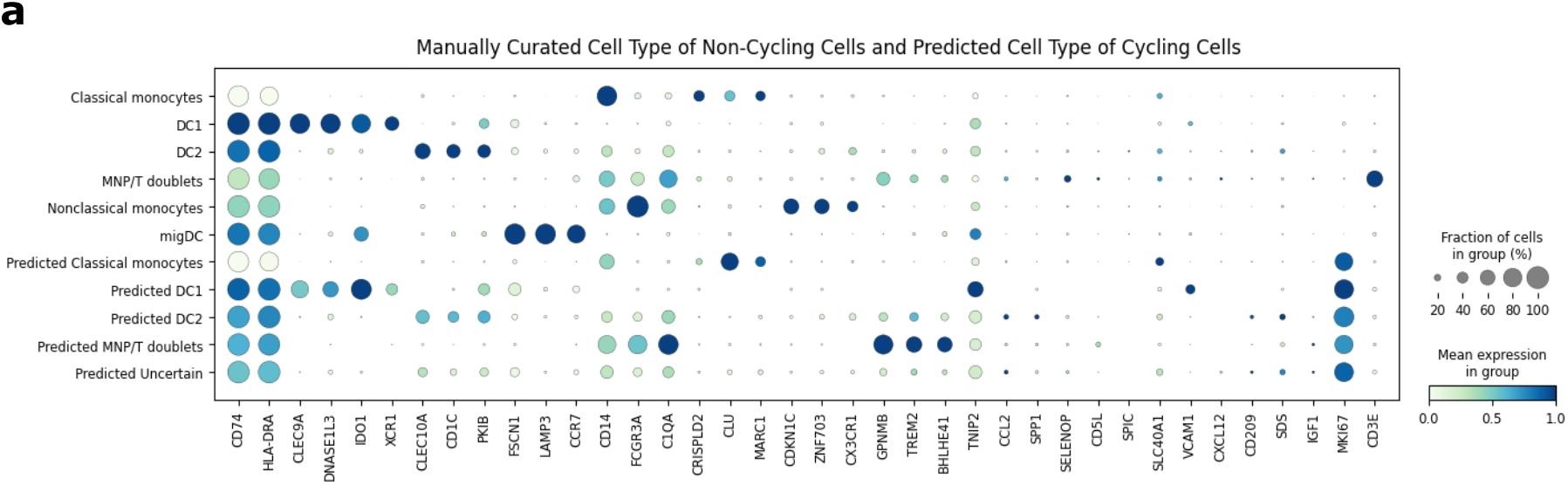
Identities of cycling cells are obtained after using CellUntangler to remove the effects of the cell cycle. (**a**) Marker gene expression across cell type of the non-cycling cells and across predicted cell type of the cycling cells. The cell type was considered uncertain if one or more of the *k*-NN classifiers predicted a different cell type from the rest.

**Supplementary Fig. 6:**
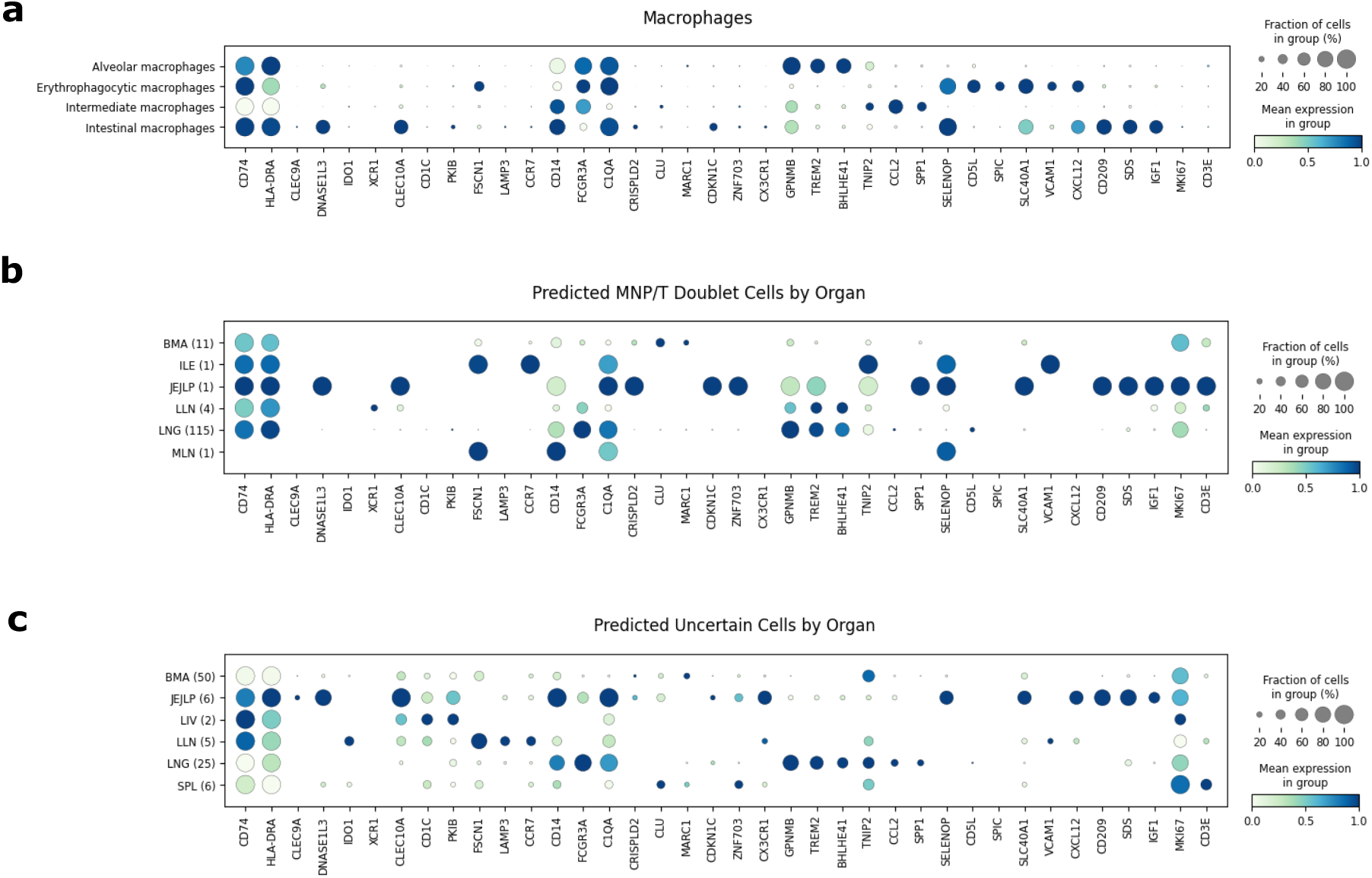
Cells predicted to be MNP/T doublets or whose identity was uncertain are likely macrophages. (**a**) Marker gene expression of the macrophages present in the original dataset of immune cells. (**b**) Marker gene expression of cycling cells predicted to be MNP/T doublets and (**c**) cycling cells whose identity was uncertain. The organs are bone marrow (BMA), ileum (ILE), the lamina propria of the jejunum (JEJLP), lung-draining lymph nodes (LLN), lung (LNG), mesenteric lymph nodes (MLN), liver (LIV), and spleen (SPL). The number of cells from each organ is indicated in the brackets. The cell type was considered uncertain if one or more of the *k*-NN classifiers predicted a different cell type from the rest.

**Supplementary Fig. 7:**
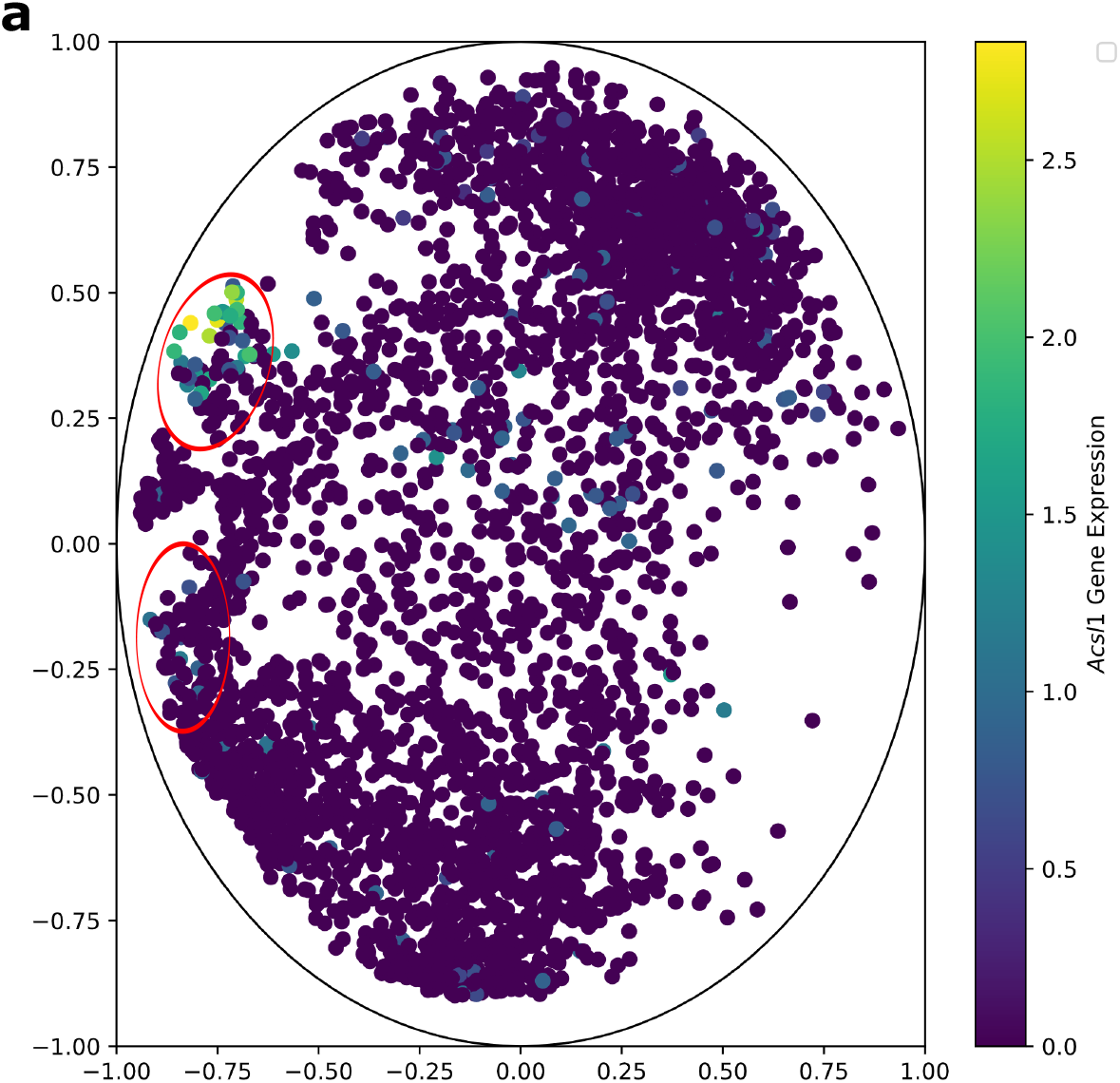
Epsilon cells are split into two groups in the second component of CellUntangler. (**a**) Cells are colored by *Acsl1* gene expression (size-factor normalized and log-transformed). The *Acsl1* ^+^ epsilon cells and *Acsl1* ^−^ epsilon cells are circled in red (above and below).

## Supplementary notes

### S1 The general CellUntangler model

We explain the general CellUntangler model. Assuming we divide the latent space into *k* subspaces, 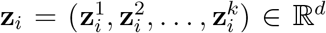. We also decompose **x**_*i*_ into *k* components, 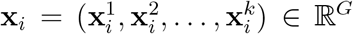, where *G* is the number of genes. Each component 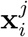 is passed to the same encoder to output the parameters for 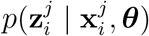. The decomposition of **x**_*i*_ will be based on which signal we want 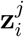 to capture. The final component, 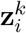, is an exception and always captures signals not accounted for by the first *k*-1 latent components. For example, assuming we have three components and we want both 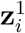 and 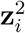 to capture the cell cycle signal then both 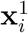 and 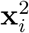 would be the UMI counts of the cell cycle marker genes. Non-cell-cycle-specific signals would be captured in 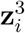. Alternatively, if we wanted 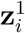 to capture the cell cycle and 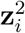 to capture cell type, 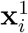 would be the UMI counts for cell cycle marker genes and 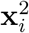 would be the UMI counts for cell type marker genes. Again, 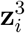 would capture any remaining signals so 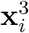 would be the UMI counts for the remaining genes which are neither cell cycle marker genes nor cell type marker genes. When decomposing **x**_*i*_, for any two components, 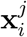 and 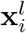, the marker genes must be either the same or disjoint. In our example of capturing the cell cycle, cell type, and remaining signals, the cell cycle marker genes would have to be disjoint from the cell type marker genes and the remaining genes.

When using the decoder to model **x**^*j*^, we leverage the latent components specifically intended to capture the intended signal. For 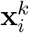, however, all latent components are utilized. The same decoder is used. For example, assuming that 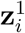 and 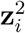 capture the cell cycle and 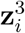 captures the non-cell cycle signals, we would model 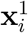 as 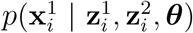. As 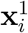 would be the same as 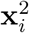, we do not model 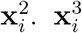 would be modeled as 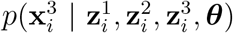. In our example of cell cycle, cell type, and remaining signals, we would model 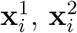, and 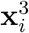 as 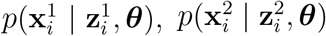, and 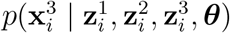.

## Supplementary tables

**Table S1:**
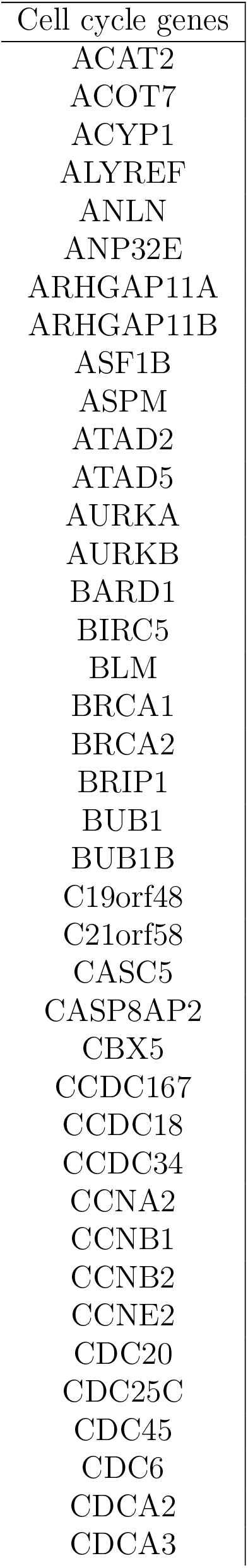
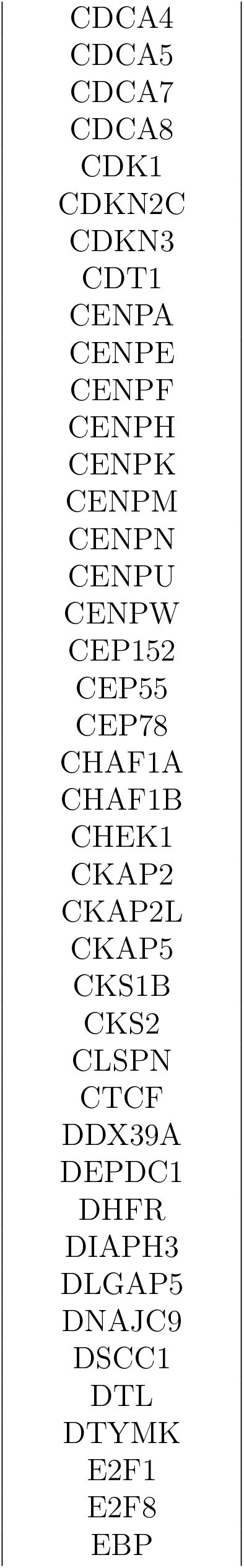
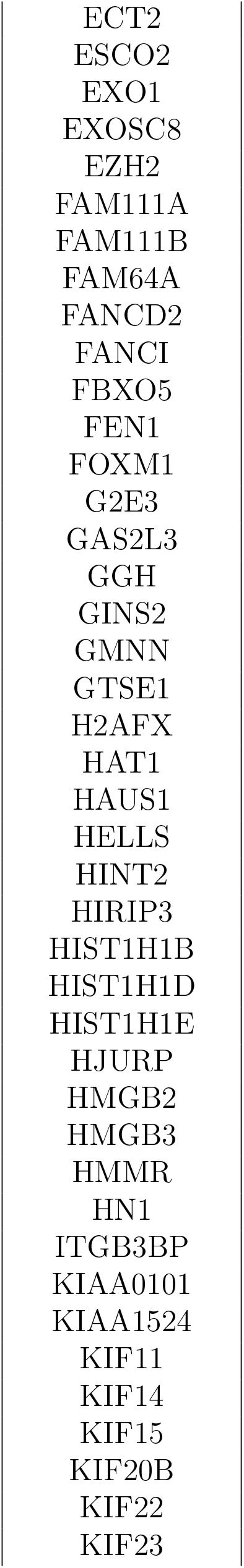
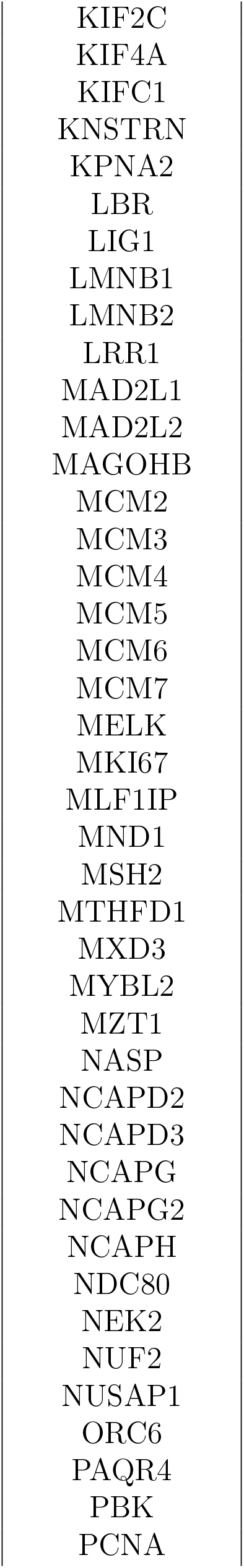
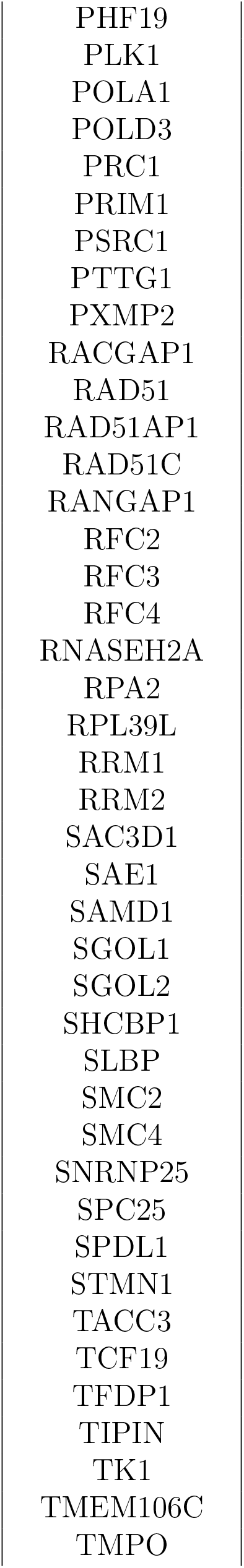
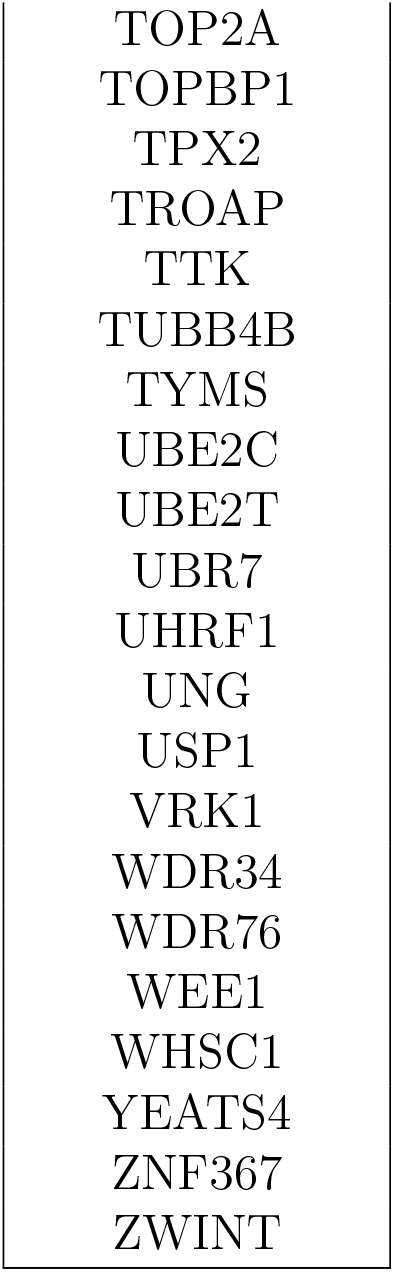
The cell cycle marker genes we used for CellUntangler.

